# The Omp85 family protein, TamA, exhibits characteristics of a suitable drug target against *Pseudomonas aeruginosa*

**DOI:** 10.1101/2025.09.16.676292

**Authors:** Rachael Duodu, David J. Boocock, Lesley Hoyles, Jack C. Leo

**Affiliations:** Antimcrobial Resistance, Omics and Microbiota Group, Centre for Systems Health and Integrated Metabolic Research, Department of Bioscience, School of Science and Technology, Nottingham Trent University, Nottingham, UK; John van Geest Cancer Research Centre, Nottingham Trent University, Nottingham, UK

**Keywords:** translocation and assembly module, outer membrane proteins, proteomics, transcriptomics

## Abstract

The outer membrane (OM) of Gram-negative bacteria is crucial for cell stability and virulence and acts as a permeability barrier. The biogenesis, assembly, and regulation of proteins in the OM are therefore attractive areas of study that could lead to identifying novel drug targets. The Translocation and Assembly Module (TAM), composed of TamA and TamB, facilitates the insertion of some β-barrel proteins into the OM of *Escherichia coli* and *Klebsiella pneumoniae*, and has also been implicated in lipid homeostasis. However, its role in *Pseudomonas aeruginosa* remains mostly uncharacterized. To investigate the TAM’s function and drug target potential in *P. aeruginosa*, we generated both single- and double-gene TAM knockouts and assessed their fitness using competition growth assays against wild-type (WT) strains. The WT significantly outcompeted the TAM mutants, indicating a fitness defect. Proteomic analysis revealed surprisingly similar profiles between WT and the double knockout strains, while single knockouts showed changes in OM proteins and reduced expression of flagellar components consistent with attenuated swimming motility observed in Δ*tamA*. Single mutants exhibited differential levels of expression of lipoproteins of the β-barrel assembly machinery suggesting compensatory OM remodelling. In vivo infection assays using *Galleria mellonella* larvae demonstrated significantly higher survival rates when infected with TAM mutants, with *tamA* mutants showing the greatest attenuation in virulence. Our findings demonstrate a role the TAM plays in *P. aeruginosa* virulence and identify TamA as a potential drug target for the development of new antimicrobial therapies.

**Data summary:** The RNAseq data reported on in this article are available from ArrayExpress: E-MTAB-15348 The mass spectrometry proteomics data have been deposited to the ProteomeXchange with the dataset identifier PXD06704.

## INTRODUCTION

The outer membrane (OM) of Gram-negative bacteria provides mechanical protection and serves as a permeability barrier due to its asymmetry, consisting of phospholipids on the inside and lipopolysaccharides (LPS) on its exterior (Funahara & Nikaido, 1980). It boasts abundant integral OM proteins (OMPs), most of which adopt a β-barrel structure.(Hermansen et al., 2022). The biogenesis of the OM involves the transport of precursors of its constituents from the cytoplasm where they are synthesized followed by insertion into the OM(Ricci & Silhavy, 2019). Nascent OMPs are transported into the periplasm through the Sec machinery. Periplasmic chaperones like SurA and Skp then escort unfolded OMPs (Chamachi et al., 2021) to the β-barrel assembly machinery (BAM) which mediates their correct folding into the OM. OMPs perform many important functions, including transport, virulence and bacterial survival, and therefore have been shown to be good drug targets (Hermansen et al., 2022).

The BAM has been extensively studied during the past decade and it is required in the assembly of most OMPs including OmpA (Hussain et al., 2020), OmpC (Hussain et al., 2021) OmpF (Lee et al., 2019) and OmpT (Hagan et al., 2010), Ag43(Rossiter et al., 2011), intimin (Bodelón et al., 2009), TolC (Malinverni et al., 2006) and PhoE (Robert et al., 2006) in *Escherichia coli*, among many others. The BAM consists of BamA, an Omp85 family integral membrane protein (Voulhoux & Tommassen, 2004;Gentle et al., 2004) connected by its 5 polypeptide transport-associated motifs (POTRA) domains to four lipoproteins, BamB, C, D and E. BamA and BamD have been demonstrated to be essential in *E. coli*.(Wu et al., 2005; Malinverni et al., 2006; Rossiter et al., 2011).

Over a decade ago, another OM insertion machinery, the translocation and assembly module (TAM), was discovered (Selkrig et al., 2012). The TAM consists of TamA, also an Omp85 family member that has structural similarity to BamA, and a second protein, TamB, which spans the periplasm and is anchored in the inner membrane (IM). TamA has three POTRA domains and interacts with TamB, which belongs to the AsmA family of proteins that were recently demonstrated to be involved in phospholipid transport in both *E. coli* (Ruiz et al., 2021) and *Pseudomonas aeruginosa* (Sposato *et al*,2024). The TAM was first thought to function in the assembly of autotransporter proteins, specifically EhaA and Ag43 of *E. coli* and p1121 from *Citrobacter rodentium* (Selkrig et al., 2012). However, the TAM only seems to be required for the assembly of a subset of autotransporters, including the inverse autotransporter intimin, whereas trimeric autotransporters are not substrates of the TAM (Heinz et al., 2016; Rooke et al., 2021). More research has since demonstrated the role of TAM in the assembly of fimbrial ushers (Josts et al., 2017 ;Stubenrauch et al., 2016) and TolC-like proteins, which form the OM channel of tripartite efflux pumps (Stubenrauch et al., 2022). Although the BAM is self-sufficient in biogenesis of fimbrial ushers and autotransporters, the absence of TAM partially affects biogenesis of autotransporters and usher proteins (Wang et al., 2024). A time-course assay was used to demonstrate efficient assembly of the usher protein FimD completed in under 2 minutes in wild-type (WT) cells, whilst this took 240 minutes in the absence of TAM (Stubenrauch et al., 2016).

Since its initial discovery (Selkrig et al.,2012) where its deletion in *Citrobacter rodentium* and *Salmonella enterica* was demonstrated to diminish virulence, TAM has been increasingly implicated in virulence in many other bacteria. TAM mutants of the fish pathogen *Edwardsiella tarda* have been shown to have reduced flagella formation with attenuated motility and invasion into host cells, and even lowered host mortality (Li et al., 2020). Furthermore, *tamB* mutant strains of *Vibrio fischeri* showed a 3.7-fold fitness defect in a competition assay (Brooks et al., 2014).

Other virulence-associated phenotypes of the TAM include causing breaches in the permeability barrier of the OM under stress conditions (Jung et al., 2021), where the deletion of *tamA* led to increased sensitivity of carbapenem resistant-*Klebsiella pneumonia* to vancomycin. The OM integrity impairment was further confirmed by a significantly higher 1-N-phenylnaphthylamine (NPN) fluorescence in *tamA* mutant in a hypo-osmotic buffer.

Apart from one recent study demonstrating an increased OM permeability and the redundancy of AsmA-like proteins including TamB in *P. aeruginosa* (Sposato et al., 2024), the role of the TAM in *P*. *aeruginosa* virulence has not been addressed *P. aeruginosa* is an opportunistic Gram-negative bacterium with a myriad of virulence factors that contribute to its success as a pathogen (Pang et al., 2019). It is notorious for being resistant to antibiotics, thereby earning a spot in the World Health Organisation’s bacterial priority pathogens list as a member of the high priority group for research on and control of antimicrobial resistance (WHO, 2024). In response to the pressing need for alternative therapeutics for the treatment of *P. aeruginosa* infection driven by the increasing emergence of antibiotic resistance (Pang et al., 2019), and considering the essential role of the OM in *P. aeruginosa* along with the established involvement of TAM in OM biogenesis; we investigated the role TAM plays in virulence in *P. aeruginosa* and analysed its suitability as a drug target against this high priority pathogen.

## MATERIALS AND METHODS

### Bacterial strains

The bacterial strains used in this study were *Escherichia coli* DH5α and SM10π*pir*. The reference laboratory strain *Pseudomonas aeruginosa* PA14 was used to generate all the knockout mutants constructed in this experiment. Bacteria were grown in lysogeny broth (LB)-Lennox at 37 °C unless otherwise stated. Room temperature (RT) is 20 °C. Antibiotics used were gentamicin (50 µg/mL) for *P. aeruginosa* or (15 µg/mL) for *E. coli*, and tetracycline (100 µg/mL) for *P. aeruginosa* unless indicated otherwise. Gene knockouts with subsequent replacement of genes with tetracycline resistance cassette were achieved by allelic exchange via homologous recombination as described by (Hmelo et al., 2015).

### Plasmid construction

Plasmid constructs used for the in-frame deletion of *tamA, tamB* and t*amAB* genes were obtained using the Platinum SuperFi II Polymerase (Thermo Fisher Scientific). Firstly, DNA fragments of approximately 1600 bp upstream and downstream of the gene of interest were amplified from PA14 genomic DNA including the start and stop codon of the genes of interest. Primers amplifying each fragment were designed to have an overhang complimentary to tetracycline resistant cassette with flanking flippase recognition target sites (FRT-tet). FRT-tet was amplified from pFRT-tet 129 (Doug Mortlock Lab:Mortlock lab unpublished plasmids) and assembled via Gibson assembly (Gibson et al., 2009) to obtain a sequence in the order upstream sequence-FRT-tet-downstream sequence in a pEXG2 vector (Hmelo et al., 2015) to obtain pEXG2 construct that was verified by colony PCR and whole-plasmid DNA Sequencing by Microbes NG. Primers used for PCR and cloning are listed in supplementary Table 1.

### Construction of *tam* knockout and Introduction of FRT-tet as a selection marker

The pEXG2 construct was introduced into *P. aeruginosa* through mobilisation using the strain SM10π*pir* by conjugation. Resultant transconjugants were selected on *Pseudomonas* isolation agar (PIA) infused with 50 μg/mL of gentamicin. Homologous recombination results with subsequent curing of the plasmid backbone using sucrose-based selection as previously described (Hmelo et al., 2015) resulted in obtaining mutants with the gene of interest replaced with the FRT-tet cassette.

### Removal of tetracycline cassette from *tam* knockouts

To remove the Tet cassette, an arabinose-inducible plasmid encoding the flp recombinase was produced. This was done by introducing a temperature-sensitive *Pseudomonas aeruginosa* origin of replication (Silo-Suh et al., 2009) into the plasmid pCMT-flp (Hossain et al., 2015). We also switched the selection marker from ampicillin to gentamicin. To increase the efficiency of the flippase expression in *P. aeruginosa*, we introduced the arabinose-inducible P_BAD_ promoter as well as the *araC* gene, amplified from the pBAD-HisA (Invitrogen) upstream of the *flp* gene to yield the plasmid pFLPGm-BAD. This was transformed into *Pseudomonas aeruginosa* knockout mutants containing the FRT-Tet cassette by preparing chemically competent cells using the TSS method (Chung et al., 1989), transforming the plasmid using heat shock and recovering at 30° C for one hour before plating onto LB with 50 µg/mL gentamicin and glucose at 0.5 %. After overnight growth at 30 °C, a transformant colony was inoculated in LB + Gm50 + glucose 0.5%. Once the culture had reached mid log phase, the medium was replaced with fresh LB + Gm50 containing 0.5 % L-arabinose, after which the culture was incubated for 3 hours to induce FLP expression. A 1:100 dilution of this culture was grown overnight on LB agar with arabinose at 0.05 % but no gentamicin at 37° C to cure the pFLPGm-BAD plasmid. A couple of colonies were inoculated in LB broth (no selection) to grow overnight at 37° C. Dilutions were plated for individual colonies the following day. Colonies were then streaked out on Gm 50 µg/mL, Tet 100 µg/mL or LB without antibiotic to look for clones sensitive to Gm and Tet. Loss of the Tet cassette and the flp plasmid was confirmed by PCR.

### Making complementation plasmids

The genes of interest were amplified from PA14 using primers binding approximately 100 bp up and downstream the genes on interest. The primers were designed such that they had overhangs complementary to pSRK-Gm (Khan et al., 2008) into which the fragment was assembled via Gibson assembly (Gibson et al., 2009) and transformed into *Escherichia coli* DH5α and plasmid DNA extracted using a plasmid miniprep kit (New England Biolabs) to obtain pSRK-Gm with a functional copy of *tamA* and *tamAB*. The construct was verified by Sanger sequencing (Source Bioscience, UK). This was then transformed into electrocompetent cells of the corresponding knockout strains that had been cured of the Tet-resistance cassette. As a control strain, pSRK-Gm was transformed into electrocompetent cells of PA14-WT to obtain pSRK-PA14WT.

### Whole-cell and OM protein extraction

OM isolation was performed following the protocol as described by Leo et al. (2015) with some adjustments. Briefly, bacteria were inoculated into 500 mL of LB and grown overnight at 37° C with shaking at 180 rpm. A number of bacteria corresponding to 500 mL at an optical density at 600 nm (OD_600nm_) of 1.0 was pelleted for 10 minutes at 5000x g. The pellet was washed with 20 mL 10 mM HEPES buffer at pH 7.4 and then resuspend in 10 mL of 10 mM HEPES, to which 0.1 mg/mL lysozyme, MgCl_2_ and MnCl_2_ at 10 mM and DNase I at 10 µg/mL were added. The cells were then lysed by sonicating on ice using a Soniprep 150 plus machine 3 x 30 seconds with one minute between pulses. The lysates were then centrifuged at 5000 × *g* for 10 minutes followed by the supernatant being transferred to an SS-34 tube and centrifuged for 30 min at 25 000 x *g*. The supernatant was decanted, and the pellet resuspended in 2 mL of 10 mM HEPES at pH 7.4 in addition to 2 mL of 2% (w/v) *N*-lauroyl sarcosine. The IMs were subsequently solubilized at room temperature (RT) with agitation for 30 min.

Following this, the tubes were centrifuged at 25 000 ×*g* for 30 min to pellet the OM followed by a 10 min wash step with 5 ml 10 mM HEPES pH 7.4 at 25000 xg. The pellet was then resuspended in 1 mL of 5% SDS with 5mM EDTA in distilled water and stored at -80 °C.

### RNA Extraction

Total RNA was extracted using Monarch total RNA Miniprep kit (New England Biolabs T2010S) following the manufacture’s extraction protocol for bacteria with some few changes. Briefly, a subculture was made from overnight cultures and grown to mid-log phase (OD_600 nm_ 0.5) at 37 °C and placed on ice immediately. Aliquots (2mL) of cultures were put into prechilled 2 mL tubes to halt cellular processes. Samples were then centrifuged in a prechilled to 4 °C centrifuge at 12 000 x g for 1min. The pellets obtained were lysed enzymatically using 100 µL of 0.1mg/mL lysozyme for 15 min and EDTA was added to 10mM. Following this step, the manufacturer’s extraction protocol was followed until RNA was eluted. Samples were then measured using the Nanodrop (Thermo Scientific NanoDrop 2000 Spectrophotometer) where samples were ensured to have A_260_/A_280_ of 1.9–2.1, and an A_260_/_230_ of 2.0 –2.2. Samples were then stored at -80 °C and transported on dry ice to Novogene (UK) for RNA-Sequencing.

### RNA sequencing and data analyses

Quality control, ribosomal RNA depletion, library preparation and sequencing (Illumina NovaSeq X) were done according to Novogene’s proprietary protocols, generating at least 6 Gbp sequence (paired end, 150 bp) data per sample.

The US web-based platform Galaxy (http://usegalaxy.org/) was used in initial analyses of transcriptomic data (Thompson et al., 2025). The genome sequence (fasta file) and annotations (gff file) for *P. aeruginosa* strain UCBPP-PA14 (acquired from https://ftp.ncbi.nlm.nih.gov/genomes/all/GCF/000/014/625/GCF_000014625.1_ASM1462v1/; 25 May 2025) were used. The quality of the sequence reads was assessed using FastQC Galaxy Version 0.74+galaxy1(Wingett & Andrews, 2018); no trimming of reads was needed as all sequence data were of high quality (Phred ≥30) and free of adapter sequences. Sequence reads were then mapped to the UCBPP-PA14 reference genome using HISAT2 (Galaxy Version 2.2.1+galaxy1) set to paired-end library, before counting the number of reads that mapped to genes using featureCounts (Galaxy v2.1.1+galaxy2). To test for differential gene expression between the Δ*tamA*, Δ*tamB* or Δ*tamAB* and WT samples, limma (Galaxy Version 3.58.1+galaxy0) (Ritchie et al., 2015) was run with the count data from featureCounts. Genes were considered significantly differentially expressed based on adjusted P value < 0.05 (Benjamini–Hochberg procedure), or P value < 0.05 and log_2_-fold change ≥1 (absolute value). Venny v2.1.0 (Oliveros, 2007-2015) was used to create a Venn diagram of the significantly differentially expressed genes. DESeq2 (Galaxy Version 2.11.40.8+galaxy0) was used to generate normalized count data for the entire dataset.

Box plots and volcano plots were generated using the R package tidyverse v2.0.0. A Kyoto Encyclopedia of Genes and Genomes (KEGG)-based network analysis was undertaken using the significantly differentially expressed genes. To do this, KEGGREST v1.46.0 was used to download information on the KEGG entry for *P. aeruginosa* strain UCBPP-PA14 (pau, https://www.genome.jp/entry/gn:T00401; accessed 29 May 2025). Nucleotide sequences of genes for the KEGG entry were downloaded and used to create a blastn database (blast v2.12.0+) against which the predicted genes of the NCBI-annotated genome used in the Galaxy-based work were searched. Hits were filtered based on the highest query coverage and identity, with this information used to map KEGG annotations to the NCBI annotations (KEGG – pau:PA14_31680 = *tamA*, pau:PA14_31690 = *tamB*; NCBI – PA14_RS12965 = *tamA*, PA14_RS12970 = *tamB*) (5889 genes mapped in total). Data were analysed using SPIA v2.58.0 (Tarca et al., 2009) and used to create a KEGG pathway-based network graph (KEGGgraph v1.66.0 (Zhang & Wiemann, 2009) igraph v2.1.4 (Csardi & Nepusz 2006) was used to generate network statistics, with RCy3 v2.26.0 (Gustavsen et al., 2019) used to export the network and visualize it with Cytoscape v3.10.3(Shannon et al., 2003).

### Mass Spectrometry

50 ug total protein was digested using trypsin following the modified S-Trap protocol (Protifi, US) as described previously (Morgan et al., 2024). Dried peptides were reconstituted in 200uL

5% acetonitrile/0.1% formic acid. Supernatants were transferred to high recovery LC vials and the autosampler kept at 8° C. Peptide separation was carried out. Peptide separation was performed on a Waters M-Class UPLC system equipped with a Phenomenex Kinetex XB-C18 column (2.6 μm, 15 × 0.3 mm) maintained at 30°C, using a linear gradient of 3–35% acetonitrile (mobile phase B) over 12 minutes at a flow rate of 10 μL/min, followed by washing at 80% B and re-equilibration to 3% B at 12 μL/min. Three microlitres of each sample were injected in direct inject mode. Mass spectrometric analysis was conducted on a Sciex 7600 ZenoTOF operating in positive ion mode with Data Independent Acquisition (DIA, zenoSWATH), employing a 25 ms TOF-MS scan and 65 variable SWATH windows (12 ms each) covering *m/z* 400–750, with a total cycle time of 1.146 s. Raw data files (.wiff, .wiff2, .wiff.scan) were processed using DIA-NN (version 1.9.2) (Demichev et al., 2020) with a *Pseudomonas aeruginosa* SwissProt FASTA database (downloaded Nov 2024), enabling library-free search and deep learning-based spectra prediction. Label-free quantification was performed at a precursor FDR of 1.0%, with default DIA-NN parameters unless otherwise specified. Data were processed using AMICA v 3.01(Didusch et al., 2022). Missing values were imputed using the minimum intensity method, and differential abundance was assessed using the limma statistical framework. Significantly changed proteins were considered at a threshold of log2fold > 2 and adjusted value p < 0.05.

### Growth curves

Overnight cultures of all strains were set up in biological quadruplicate in LB (gentamicin was added at 50 µg/mL to complemented strains). As a control, pSRK-PA14WT was used in comparisons with the complemented strains. The turbidity of the cultures was measured and adjusted to an OD_600nm_ of 0.1. 2 µL of diluted cultures was added to 200 µL of LB (with gentamicin at 50 µg/mL for complemented strains) in wells of a 96-well plate and sealed with a Breath-Easy membrane (Sigma-Aldrich). 200 µL of LB was inoculated into separate wells as a sterility control. The plate was then incubated at 37° C in a Cytation 3 plate reader with shaking and OD_600_ was measured every 20 minutes for 24 hours.

### Competition assay Growth competition of *tam* mutant versus knockouts

Mutant strains containing the tet cassette insert and PA14-WT were grown on *Pseudomonas* isolation agar (PIA) with 100 μg/mL of tetracycline and just PIA, respectively. Single colonies were inoculated into 5 mL of LB broth and grown overnight 37 °C with 100 μg/mL of tetracycline for mutant strains. This was followed by measuring cultures and adjusting with LB to an OD_600nm_ of 1. Mutant and PA14 were inoculated into LB broth in a ratio 1:1. Before the mixed culture was incubated, it was inoculated onto PIA-tet 100 and PIA only to ascertain that both strains were present at equal amounts and viable. The mixed cultures were retrieved the following day and serially diluted and plated for single colonies. Each culture was plated both on PIA only and on PIA-tet. Colonies were counted the following day, where the WT numbers were obtained by subtracting colonies on PIA-tet (mutant with tetracycline cassette) from total colonies on PIA only (both mutants with tet cassette and wildtype).

### Biofilm formation

Biofilm assay was performed following the protocol as described by (Coffey & Anderson, 2014) with incorporated adjustment from protocol of Merritt et al. (2005).

Bacteria were inoculated into 5 mL of LB broth and grown overnight at 37 °C with shaking. Cultures were adjusted to an OD_600_ of 1 the following day and then diluted 1:100. Following this, 100 μL of the diluted cultures were aliquoted into 96-well microtiter plate (Sigma-Aldrich, Costar) and incubated at 37 °C for 18 hours. After this, the cultures were aspirated using a 1 mL pipette and washed 3 times with 200 μL of PBS to remove non-adherent cells. Excess PBS was removed by patting the plate firmly on a paper towel. The plate was then incubated for 1 hour at 60 °C to fix adherent cells. Next, the biofilm was stained by adding 150 µL 1 % (w/v) crystal violet followed by a 20-minute incubation at RT. Plates were rinsed three times with distilled water and inverted to dry. Finally, 150 µL of 33 % (v/v) glacial acetic acid was added to each well to solubilize the stained biofilm, after which the supernatants were transferred to a new plate and the OD_540_ was measured using a Cytation 3 plate reader.

### Swimming motility

Swimming motility assays were performed following the protocol of (Ha Dae-Gon et al., 2014). 5× M8 solution was prepared with 64 g Na2HPO4·7H2O, 15 g KH2PO4: 2.5 g NaCl in 1 litre of water. 200 mL of 5× M8 solution was added to 800 mL of 0.3 % autoclaved agar to yield a homogenous suspension. Glucose or casamino acid were added to a concentration 0.2 % and 0.5 %, respectively. MgSO_4_ was then added to 1 mM and mixed thoroughly, and 25 mL of the mixture was poured into plates to solidify at RT. A sterile 10 µL pipette tip was dipped into an overnight culture of strains without the tetracycline cassette (Tet-sensitive strains) and PA14-WT and stabbed into the agar layer of the plate carefully to avoid touching the base of the plate. Plates were incubated upright for 18 hrs and the diameter of the cultures was measured.

### Antibiotic susceptibility test

Disk diffusion susceptibility testing was performed in accordance with the recommendations of EUCAST as described by Matuschek et al. (2014). Cartridges of antibiotic-containing disks were stored at -20°C and allowed to equilibrate to room temperature prior to use. Overnight cultures of bacteria (quadruplicates for each strain) were obtained by inoculating a single colony into 5 mL of Mueller-Hinton broth (MHB) and diluted to OD_600nm_ at 0.1 the next day. Mueller-Hinton agar plates of thickness 20 mm were inoculated with the diluted bacteria using a sterile cotton swab and disks were placed onto the inoculated media within 3 minutes of inoculation. Plates were incubated at 37 °C for 24 hrs and the diameter of the zones of inhibitions measured.

### Chelator and detergent assay

Samples were prepared in the same way as for antibiotic susceptibility testing detailed above. Sterile empty 6 mm disks were soaked with 10 µl of 20mM EDTA or 5 µl of 10 % SDS and 10 µL of 20mM EDTA.

### OM permeability testing

OM permeability testing was done using NPN following the procedure described in (Helander & Mattila-Sandholm, 2000). Briefly, in a black microtiter plate, 50 µL of 40 µM NPN in 5 mM HEPES at pH 7.2 was aliquoted into wells and topped up with 50 µL of EDTA at 20 mM. 100 µL of bacterial suspension adjusted to an OD_600nm_ value of 0.1 in 5 mM HEPES was added, and fluorescence measured at 350 and 420 nm excitation and emission respectively within 3 min in a Cytation 3 plate reader. Control wells were prepared as follows; (a) 200 µL HEPES only (b) HEPES (150 µL) plus NPN (50 µL) (c) bacterial suspension (100 µL) and HEPES (100 µL) and lastly (d) bacterial suspension (100 µL), NPN (50 µL) and HEPES (50 µL).

Results were expressed as relative fluorescence units (RFU), which were calculated as the fluorescence value of the cell suspension and NPN without the test substance subtracted from the corresponding value of the cell suspension with EDTA and NPN.

### In vivo testing of *tam* knockouts using *Galleria mellonella*

We used *Galleria mellonella* larvae (purchased from the *Galleria mellonella* Research Centre, University of Exeter, Exeter, UK) for infection assays. Mutant strains without tetracycline cassette insert or PA14-WT were cultured overnight at 37 °C and subcultures were made the following day to grow until an OD_600_ of 0.4–0.5. Following this, cultures were diluted to specific CFU/mL in PBS, based on a standard curve relating CFU/mL to OD_600nm_. PBS was used as a negative control while heat-killed PA14 (20 min incubation at 80 °C) served as control for testing the toxicity of bacterial cell components. *Galleria* injections were done using 1 ml syringes with 27 G needles with a mechanical syringe pump (Cole-Parmer, St Neots, UK) at a rate of 13.21 mL/h and a volume of 10 µL per injection. Injections were made at the bottom left proleg into the hemocoel of the larvae. To verify the inoculum doses, 10 µL of inocula were diluted and plated. The health score of larvae was calculated following the scale of (Serrano et al., 2023). The health of larvae is directly proportional to the heath score whereby higher scores reflect healthy larvae.

## RESULTS

### The *tam* mutants and their complemented strains

We first generated *tam* mutants in the *P. aeruginosa* PA14 background by standard marker exchange mutagenesis, where the *tamA* or *tamB* or the whole operon *tamAB* were replaced with a tetracycline resistance cassette flanked by FRT sites for later removal using the Flp recombinase (Hoang et al., 1998;Ishikawa & Hori, 2013). This approach was utilized to enable us to perform competition assays and select for mutants using tetracycline. The resultant mutants are referred to as Δ*tamA-tetR,* Δ*tamB-tetR,* Δ*tamAB-tetR.* Removal of the tetracycline cassette with a novel Flp-encoding plasmid that replicates conditionally in *P. aeruginosa* resulted in mutants which were sensitive to tetracycline, namely Δ*tamA,* Δ*tamB* and Δ*tamAB* (Supplementary Figure 1A).

Complemented strains with the genes *tamA* or *tamAB* cloned into the pSRKGm plasmid background (Khan et al., 2008) were dubbed Δ*tamA*::pSRK-*tamA* and Δ*tamAB*::pSRK-*tamAB,* respectively. We were unable to complement Δ*tamB*, despite several attempts. Unsuccessful attempts were also made by Ramezanifard et al. (2023) to complement *tam* single mutants, *tamA* or *tamB,* in *Salmonella.* This could be due to the *tamA* stop codon overlapping with the start codon of *tamB* (Supplementary Figure 1B), leading to difficulties in expression or the construct being toxic. We therefore complemented Δ*tamB* with the pSRK-*tamAB*.

### Growth rates of *tam* mutants and the WT are similar, but mutants are outcompeted in a mixed culture

We Compared the growth rates of *tam* knockout strains and the WT by following growth over 24- hours and generating growth curves (Supplementary Figure 2). Among all the strains, the Δ*tamA* mutant exhibited the fastest growth rate. The other strains tested all showed growth rates similar to the WT. In a competition growth assay, however, where we mixed an equal inoculum of the WT and mutant, the *tam* mutants were significantly outcompeted by the WT (PA14-WT) (Figure. 1). Moreover, the competitive index was partially increased or fully restored when complemented with *tamA* or *tamAB*, respectively. However, complementation of the Δ*tamB* with pSRK-*tamAB* did not improve their competitive index in any way, possibly because of the presence of an additional copy of *tamA*.

**Figure 1.**
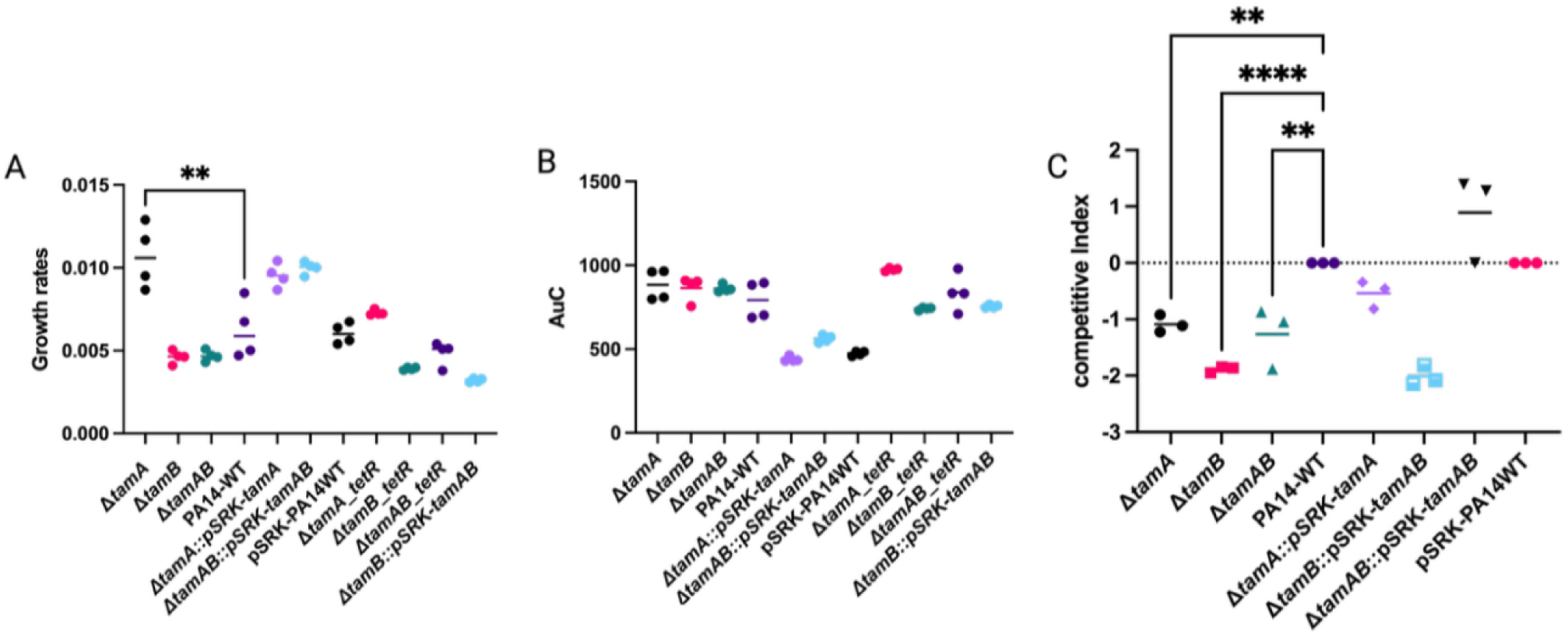
Growth and fitness of *tam* knockouts. (A) Growth rates of *tam* knockouts (Δ*tamA,* Δt*amB* and Δ*tamAB)*, complemented strains (Δ*tamA*::pSRK-*tamA*, Δ*tamB*::pSRK-*tamAB* and Δ*tamAB*::psRK-*tamAB*) and tetracycline resistant strains (Δ*tamA-*tetR, Δ*tamB*-tetR, and Δ*tamAB*-tetR). PA14-WT refers to the WT and the pSRK-PA14WT refers to the WT with an empty pSRKGm plasmid. (B) Area under the Curve (AuC) reflecting the total growth of strains. Growth was obtained using LB broth at 37° C for 24-hr growth (n=4) and growth rates and area under curve obtained using the *growthcurver* package in R (Sprouffske & Wagner, 2016) from the growth curves in (supplementary figure 2). Statistical analysis was done for both panels A and B using one-way ANOVA with Dunnett’s post-hoc test; **p < 0.005 compared to the WT (PA14-WT). (B) Growth competition between *tam* knockouts or complemented strains vs WT _g_rowth was performed in LB a 37° C by growing equal amounts of each mutant and WT together. The bars indicate competitive median. Competitive index was calculated as the ratio of CFU of mutant to WT. Values below and above 0 indicates growth defect or growth advantage differential to the WT respectively. Statistical analysis used was one way ANOVA with Dunnett’s post-hoc test; **p < 0.01, **** p < 0.0001.

### Deletion of *tamA* or *tamB* differentially impacts the OM proteome

To verify whether deletions of *tamA, tamB* or *tamAB* impact the proteome of the mutants, we extracted the whole-cell protein (WCP) and the outer membrane proteins (OMP) of the mutants and WT and analysed their proteome via mass spectrometry. In the OMP samples, we observed many significantly differentially expressed proteins between the single mutants compared to the WT (Figure 2A and B). However, many of these were cytosolic proteins, which we assume are spurious contaminants in the OMP samples. We therefore manually filtered for cell envelope proteins (Table 1) leaving out all the cytoplasmic proteins (Supplementary Table 2 & 3) from our analysis, especially because these cytoplasmic proteins were not significantly differentially expressed in the proteome analysis of the WCP (Figure 2 D & E). Proteomic profiling of the single mutants exhibited significant changes in proteins related to OM structure, transport and motility with some few differences between each mutant. In Δ*tamA*, differential expression was dominated by OM-associated proteins including TonB-dependent receptors, TolC-like proteins, the OM assembly factor BamB, and porins. Also, CheV and flagellar components found in their proteome suggests modulation of chemotaxis and motility pathways. There were many other flagellar components differentially expressed which did not reach significant levels (Supplementary Table 4), however they possibly contributed to the negative impact of swimming motility in Δ*tamA*.

**Figure 2.**
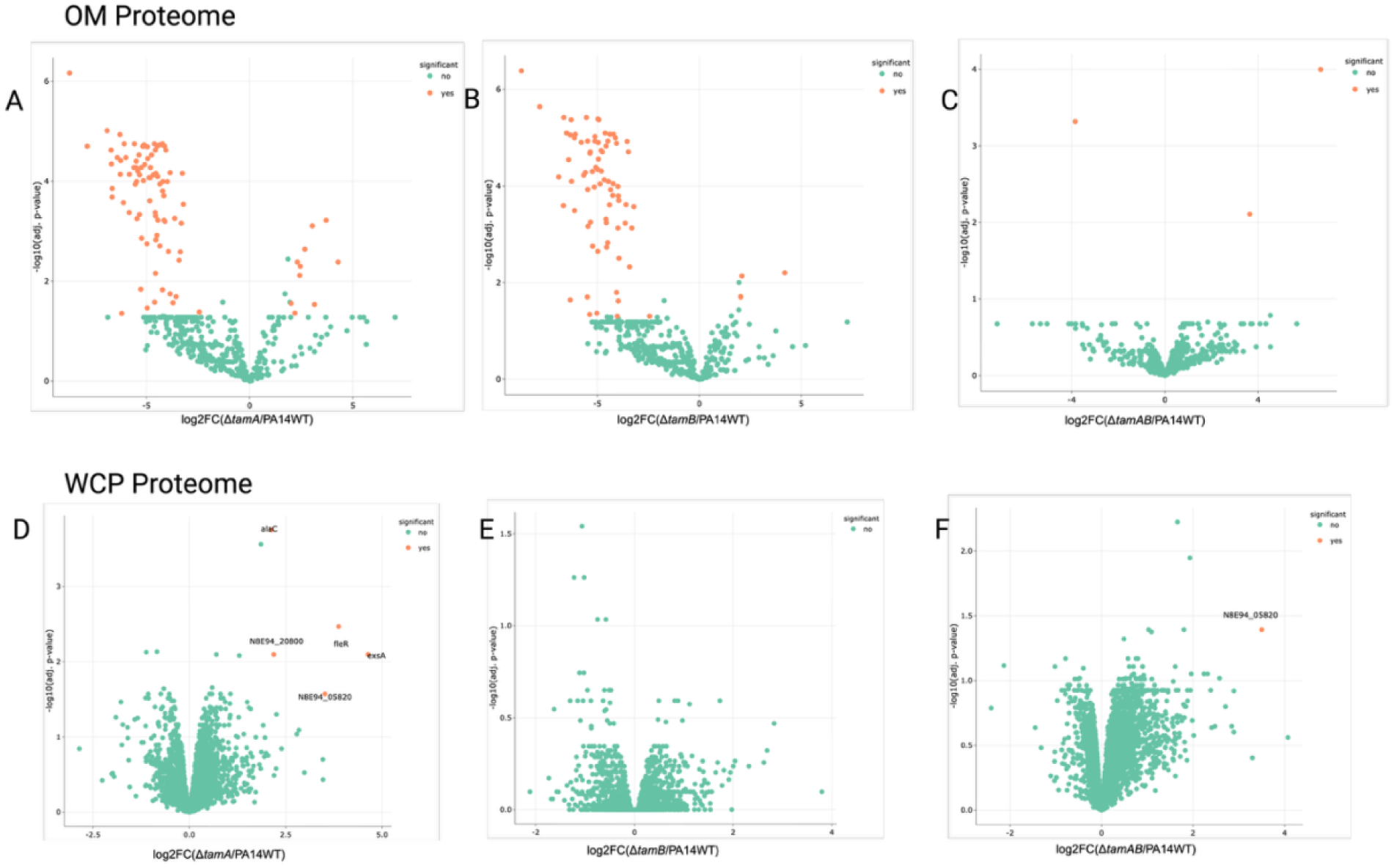
Differential envelope or whole cell protein abundances in whole cell proteins or outer membrane samples respectively of *tam* knockouts. Volcano plots showing the results of proteomic analysis of mutants and WT of: (A-C) outer membrane protein extracts (D-F) whole cell protein extracts. Orange dots represent proteins significantly upregulated or downregulated. Differentially abundant proteins identified by label-free quantification (LFQ) using DIA-NN and analysed with *amica* (https://bioapps.maxperutzlabs.ac.at/app/amica). Proteins were filtered to retain those with at least two razor/unique peptides and a minimum of three MS/MS counts in at least one group. Missing values were imputed using the minimum intensity method, and differential abundance was assessed using the limma statistical framework. Significantly changed proteins were considered from log2fold > 2 and adjusted value p < 0.05.

**Table 1.** Outer membrane proteins (OMP) expressed at significant levels in *Δ*t*amA* and *Δ*t*amB* mutant proteome compared with WT.

| *ΔtamA* | UNIPROT PROTEIN ID | PROTEIN DESCRIPTION | logFC |
| --- | --- | --- | --- |
|  | A0A7M2ZZQ8_PSEAI | Channel protein TolC | -6.287048852 |
|  | A0A0H2Z834_PSEAB | Flagellar hook-associated protein 2 | -5.475610636 |
|  | A0A0H2ZDT9_PSEAB | Chemotaxis protein CheV | -5.362046911 |
|  | A6V673_PSEA7 | Porin | -5.122999336 |
|  | A0A0H2ZH83_PSEAB | TolC family protein | -5.117572791 |
|  | A0A0H2ZGG4_PSEAB | Putative lipoprotein | -4.953177921 |
|  | A0A0H2Z6U8_PSEAB | TonB-dependent receptor | -4.86532359 |
|  | A0A0H2ZHF8_PSEAB | Putative outer membrane protein | -2.439195285 |
|  | A0A0H2ZJN1_PSEAB | Outer membrane assembly lipoprotein YfiO | 2.421652005 |
|  | YFIB_PSEAE | Outer-membrane lipoprotein YfiB | 3.03583481 |
|  | Q9HXJ7 | Outer membrane protein assembly factor BamB | 4.282713113 |
| *ΔtamB* |  |  |  |
|  | A0A2R4B1G8_PSEAI | Putative porin | -7.824455707 |
|  | A0A643IRS3_PSEAI | Porin | -6.642985962 |
|  | A0A0H2ZA96_PSEAB | Transporter | -6.511634235 |
|  | A0A0H2Z917_PSEAB | Putative purine-binding chemotaxis protein CheW | -6.326043707 |
|  | A0A7M2ZZQ8_PSEAI | Channel protein TolC | -6.287048852 |
|  | A0A9Q9N8P1_PSEAI | Putative secretion system protein | -6.123551896 |
|  | A0A0H2Z834_PSEAB | Flagellar hook-associated protein 2 | -5.475610636 |
|  | A0A0H2ZDT9_PSEAB | Chemotaxis protein CheV | -5.362046911 |
|  | A6V673_PSEA7 | Porin | -5.122999336 |
|  | A0A072ZBY7_PSEAI | TolC family protein | -5.117572791 |
|  | A0A0H2ZCQ2_PSEAB | Putative TonB-dependent receptor | -4.967871431 |
|  | A0A0H2ZFL2_PSEAB | Putative outer membrane protein | -4.209489762 |
|  | PILY1_PSEAB | Type IV pilus biogenesis factor PilY1 | -4.058229298 |
|  | A0A0H2Z825_PSEAB | Flagellar basal-body rod protein FlgG | -3.94962816 |
|  | A0A0H2ZGA2_PSEAB | RND efflux membrane fusion protein | -3.313169301 |
|  | BAMB_PSEAE | Outer membrane protein assembly factor BamB | 1.941419858 |
|  | A6V2I3_PSEA7 | Lipoprotein, putative | 2.027421614 |

In addition to the shared components like a CheV, porins and TolC, *tamB* mutants showed an expanded set of differentially expressed proteins including CheW, a coupling protein that connects CheA to receptors to facilitate chemotaxis (Szurmant & Ordal, 2004); FlgG which forms the distal rod as a drive shaft to transmit torque from the motor to the filament (Fujii et al., 2017) and PilY1, a type IV pilus biogenesis factor which mediates twitching motility (Burrows, 2012). Also, the Δ*tamB* proteome revealed a reduction of a putative secretion system protein and an RND efflux membrane fusion protein, suggesting a broader change in secretion and efflux systems.

There were surprisingly few differentially expressed proteins in the OMP proteome, however, when *tamAB* was deleted. The only three significantly different proteins were all cytoplasmic proteins namely transketolase, DNA topoisomerase IV and DUF2599 domain-containing protein, all of which were not considered in our analysis as they were not differentially significantly expressed in the WCP.

In general, the WCP samples did not highlight many significant differences (Figure 2 D-F). It is worth noting that membrane proteins (including OMPs) are only a minor component of the WCP and are likely largely missed in these samples. For Δ*tamA* WCP, however, there were 5 differentially increased expression AlaC, N8E94_20800, N8E94_05820, FleR, ExsA, indicating an up-regulation in components involved metabolic pathways, transport systems, motility and secretion systems. Δ*tamAB* had only one significant differentially upregulated protein, N8E94_05820, an uncharacterized RDD family membrane protein in *P. aeruginosa*. The RDD family constitutes a category of proteins characterized as a novel Na^+^(Li^+^,K^+^)/H^+^ antiporter in *Halobacillus andaensis* (Shao et al., 2018). Na^+^(Li^+^,K^+^)/H^+^ antiporters maintains cellular homeostasis (Meng et al., 2017). There were no differentially expressed significant proteins in the WCP of Δ*tamB.* These findings highlight the minimal (Δ*tamA)* to null significant (Δ*tamB* or Δ*tamAB)* effect the *tam* has on the cytoplasmic protein profile of *P. aeruginosa* when deleted as opposed to the impact it has on outer membrane proteome when the single genes are deleted.

### Deletion of *tam* negatively impacts biofilm formation abilities of mutants

Since there were no obvious differences in the proteome of the WCP of *tam* mutants, we decided to probe further by performing transcriptomics analysis to investigate differences, if any, at the mRNA level.

Limma-based analyses of the transcriptomic data using an adjusted P value threshold of 0.05 (Benjamini–Hochberg) showed that in the Δ*tamA* mutant, only the expression of *tamA* was significantly downregulated (P = 6.55×10^-4^) (log_2_fold change = -7.03927821) compared with the WT, in the *tamB* mutant only the expression of *tamB* was significantly downregulated (P = 4.77×10^-5^) (log_2_fold change = -7.54596815), and in the *tamAB* mutant the expression of both *tamA* and *tamB* was significantly downregulated (P = 4.72×10^-4^ and = 5.79×10^-5^, respectively) (log_2_fold changes -6.95671808 and -6.98747073, respectively) (Figure 3A), demonstrating that the knockouts had been successful.

**Figure 3.**
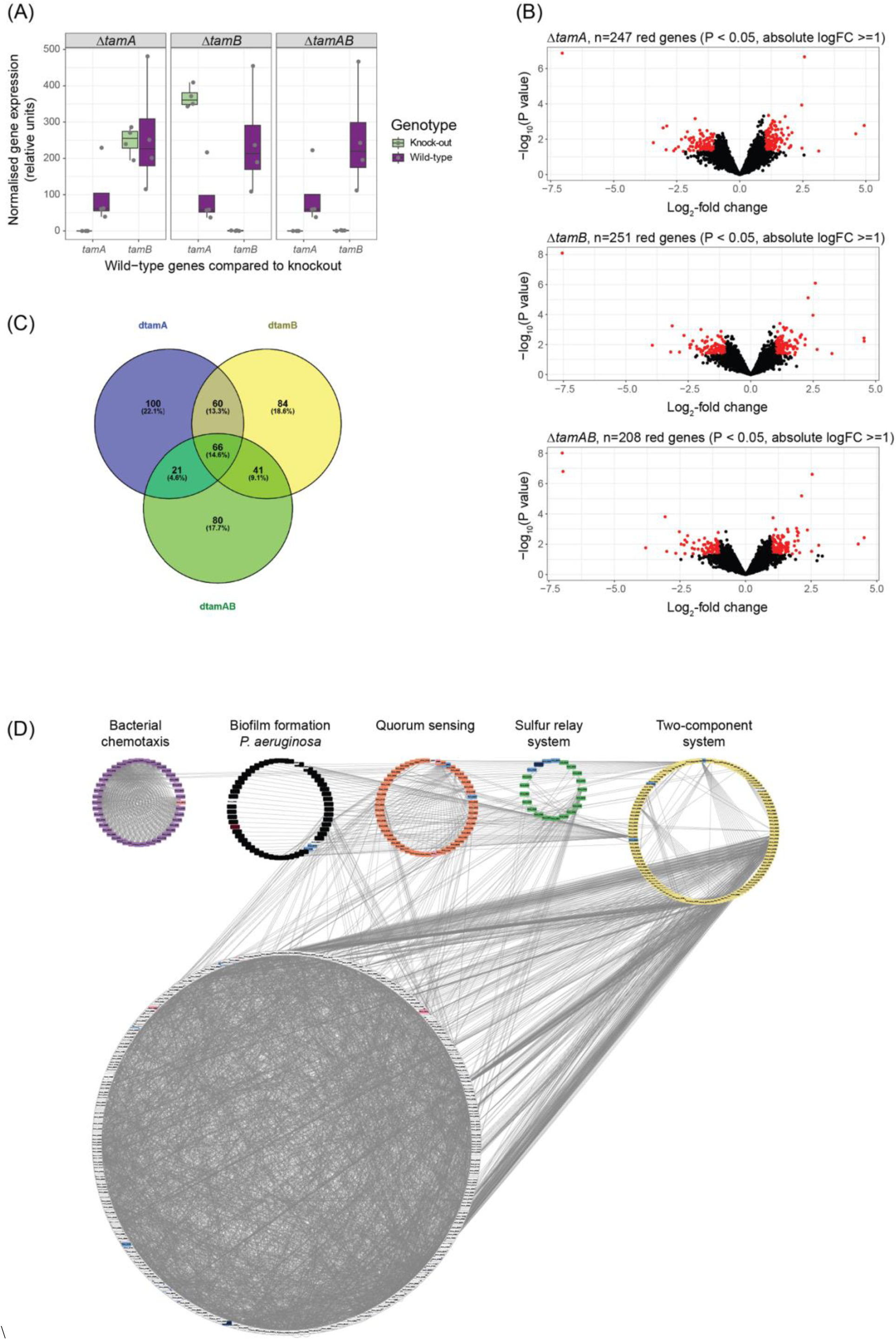
Analyses of transcriptomic data for the Δ*tamA*, Δ*tamB* and Δ*tamAB* mutants compared with the WT strain. (A) DESeq2-normalised data were plotted for the single- and double-mutant strains compared with the WT, confirming abrogation of gene expression in the mutants. n=4 biological replicates per strain. (B) Volcano plots of log_2_-transformed normalised gene expression data. Significantly differentially expressed genes (P < 0.05, absolute log_2_-fold change ≥1) are shown in red; all other genes are shown in black. (C) The significantly differentially expressed genes from (B) were compared and represented in a Venn diagram. (D) A KEGG-based pathway analysis was carried out using genome data from *P. aeruginosa* strain UCBPP-PA14. The significantly differentially expressed genes identified in the comparison of the Δ*tamA* and WT data are shown in blue and red (the darker each colour the lower or higher the log_2_-fold change in gene expression compared with the WT, respectively). The five pathways identified by SPIA as being inhibited by disruption of *tamA* are highlighted in the circular visualisation of the KEGG pathway (network) data.

Using a less conservative approach to analyses of the data (unadjusted P value < 0.05 and absolute log_2_-fold change ≥1), in Δ*tamA* 247 genes (229 with KEGG annotations) were significantly differentially expressed, in Δ*tamB* 251 genes (238 with KEGG annotations) were significantly differentially expressed, and in Δ*tamAB* 208 genes (196 with KEGG annotations) were significantly differentially expressed compared with the WT (Figure 3B). Sixty-six significantly differentially expressed genes were shared by the single- and double-knockout mutants (Figure 3C).

SPIA was used to map the significantly differentially expressed genes and their log_2_-fold change data onto KEGG pathways, to determine whether any of the pathways encoded by *P. aeruginosa* were potentially activated or inhibited by the gene mutations. At an adjusted P value threshold of 0.05 (Benjamini–Hochberg), the sulfur relay system was significantly inhibited (P = 0.047) in the Δ*tamA* mutant (Supplementary Table 5). The expression of the *tusBCDE* genes associated with this pathway (http://www.genome.jp/dbget-bin/show_pathway?pau04122+PA14_30380+PA14_30390+PA14_30400+PA14_30370) was significantly downregulated (Figure 3D). Other pathways predicted to be inhibited in the Δ*tamA* mutant were biofilm formation (in agreement with the phenotypic data below), quorum sensing, two-component system and bacterial chemotaxis. These same pathways were predicted to be inhibited in the Δ*tamB* and Δ*tamAB* mutants. SPIA of the 58/66 significantly differentially expressed genes with KEGG annotations shared by the Δ*tamA*, Δ*tamB* and Δ*tamAB* mutants showed all the aforementioned pathways except quorum sensing were associated with these genes (Supplementary Table 5).

### The OM integrity of mutants is impaired

The integrity of the OM of mutants was assessed using NPN, which is a hydrophobic compound that fluoresces when it accesses the hydrophobic interior of the OM (Helander & Mattila-Sandholm, 2000). This only happens when the barrier function of the OM is impaired. Upon treatment of *tam* mutants or wild type cells with EDTA followed by NPN, Δ*tamB* cells showed the highest fluorescence, followed by Δ*tamA* and Δ*tamAB,* all significantly higher than the fluorescence in the wild type cells showing the mutants’ OMs were more perturbed by EDTA and OM impairment among the *tam* mutants. (Figure 4A).

**Figure 4.**
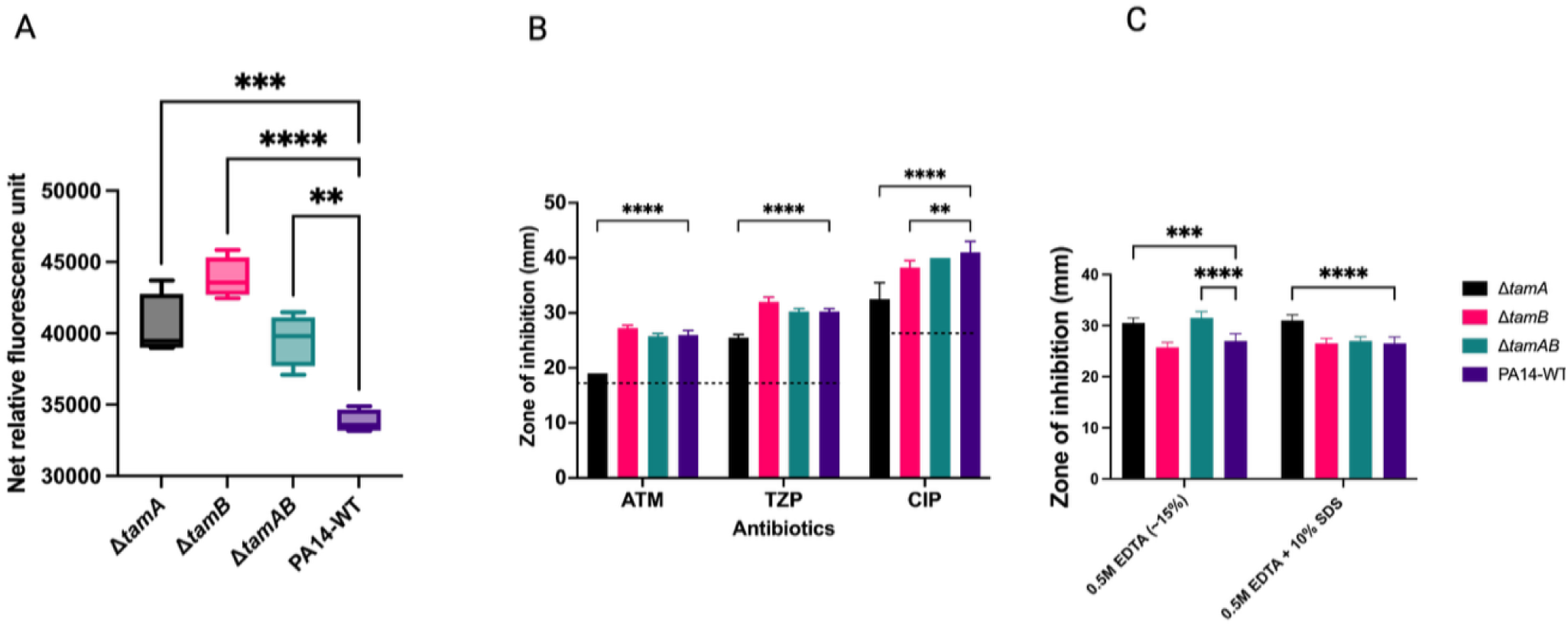
Assessing the OM integrity of *tam* mutants. **(A)** Oms of mutants are more permeable than the WT in the presence of EDTA. Permeability of the OM was assessed using an NPN assay (Helander & Mattila-Sandholm, 2000). Relative fluorescence units were calculated as the fluorescence value of the bacterial cell suspension and NPN without the test substance (no EDTA) subtracted from the corresponding value of the cell suspension with EDTA and NPN. **P<0.05; ***P<0.001; ****P<0.0001 determined by One-way ANOVA with Dunnett’s post-test. **(B**) Antibiotic susceptibility testing *P. aeruginosa* PA14 wildtype and TAM knockout using disk diffusion assays. Antibiotics Aztreonam (ATM), tazobactam-piperacillin (TZP) and Ciprofloxacin (CIP) were used. Disk diffusion assay was performed and zones of inhibition measured in mm (n =4). Black horizontal dashes across the graph represent EUCAST clinical breakpoints for resistance (EUCAST, 2025). **p<0.01; ****p <0.0001 **(C)**. Effect of chelators and detergents as measured by disk diffusion assays. The combination of EDTA and SDS enhances antimicrobial activity against Δ*tamA*. ***p<0.01; ****p <0.0001. Statistical analysis for both graph **B** and **C** determined by 2-way ANOVA with Dunnett’s post-test.

We also tested the effect of several antibiotics used to treat *P. aeruginosa* infections, as an impaired OM barrier could increase the sensitivity of the mutants. The antibiotics levofloxacin, tobramycin, meropenem, polymyxin B and colistin showed no increased efficacy in the mutants compared to the WT (Supplementary Figure 4); however, for aztreonam (ATM) and tazobactam-piperacillin (TZP), the efficacy against the Δ*tamA* mutant was reduced, though still above the EUCAST clinical breakpoints (EUCAST, 2025). Similarly, ciprofloxacin had significantly reduced efficacy against Δ*tamA* and Δ*tamB* (Figure 4B). We also tested effect of the chelator EDTA and the ionic detergent SDS and found that Δ*tamA* and Δ*tamAB* were significantly more susceptible to EDTA compared to the WT. The combination of EDTA and SDS enhanced antimicrobial activity against Δ*tamA*.

### Virulence markers in *tam* knockouts are attenuated

As *P. aeruginosa* is infamous for forming biofilm, we tested the mutants’ ability to form biofilm. Biofilm formation was reduced significantly in all mutants (Figure 5A). Biofilm formation was partially restored in complemented strains as seen by an increase in biomass, although this did not reach statistical significance (Supplementary Figure 5A).

**Figure 5.**
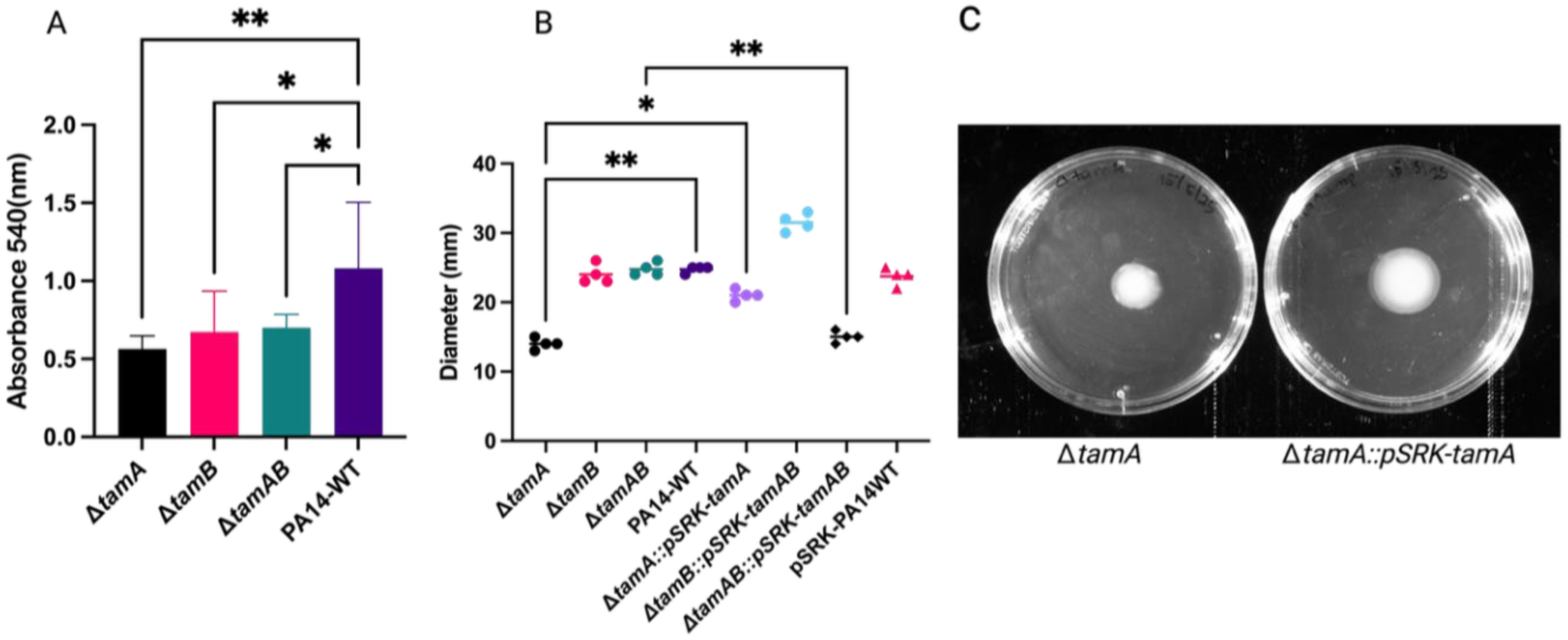
Impact of *tam* deletion on biofilm formation and swimming motility. (A) Biofilm formation of *P. aeruginosa* PA14 WT versus *tam* knockouts. 96 well microtiter plates were inoculated with PA14-WT or *tam* mutants and biofilm formation quantified using the crystal violet assay. Absorbance was measured at 540 nm. Bars show the means of 6 biological replicates with error bars denoting standard deviations. p <0.05 and **p= 0.001(p< 0.005). (B) Swimming motility in semi-solid agar showing 4 biological replicates of each strain with lines depicting the median value. (C) **p<0.005; *p< 0.05. Statistical analysis obtained using one way ANOVA with Dunnett’s post-test. Non-significant (ns) differences not displayed. (C) An image showing the swimming diameter of Δ*tamA* and its complemented strain Δ*tamA*::pSRK-*tamA*.

We proceeded to investigate whether swimming motility was attenuated since we observed a reduction in the flagellar components in the mutants in the proteomic analysis. Swimming motility of Δ*tamA* was impaired significantly when assessed in a semi-solid agar motility assay, and this was significantly restored when complemented (Figure 5B & C). The other mutants, Δ*tamB* and Δ*tamAB*, showed no reduction in their swimming abilities; however, when Δ*tamB* was complemented with *tamAB* in a pSRK background, this resulted in improved swimming motility recording the largest diameter (although not significantly) among all strains suggesting an extra copy of *tamA* might cause an additive improvement in Δ*tamB*::pSRK-*tamAB.* (Figure 5B). We also tested twitching motility, which is mediated by type 4 pili. However, no significant differences were observed between the *tam* mutants and WT (Supplementary Figure 5B).

### Reduced pathogenicity of *tamA* knockouts in vivo

To investigate whether the TAM plays a major role in *P. aeruginosa* virulence, we tested the virulence of the *tam* knockouts in a *Galleria mellonella* larval infection model (Serrano et al., 2023) Only *tamA* mutants were seen to be significantly less pathogenic as infected larvae experienced reduced mortality (Figure 6). The overall health score (Serano et al, 2023) of all the *tam* mutants were significantly higher after 24 hours compared with the WT, even though the larvae died: the larvae were not as melanized when infected with the mutants, suggesting a slight reduction in virulence. The WT PA14 not only killed the *Galleria* within 24 hours, but the tissue was also largely disintegrated, again suggesting the WT bacteria were more virulent. The negative controls (heat-killed bacteria and PBS) did not result in any mortality or significant reduction in health score.

**Figure 6.**
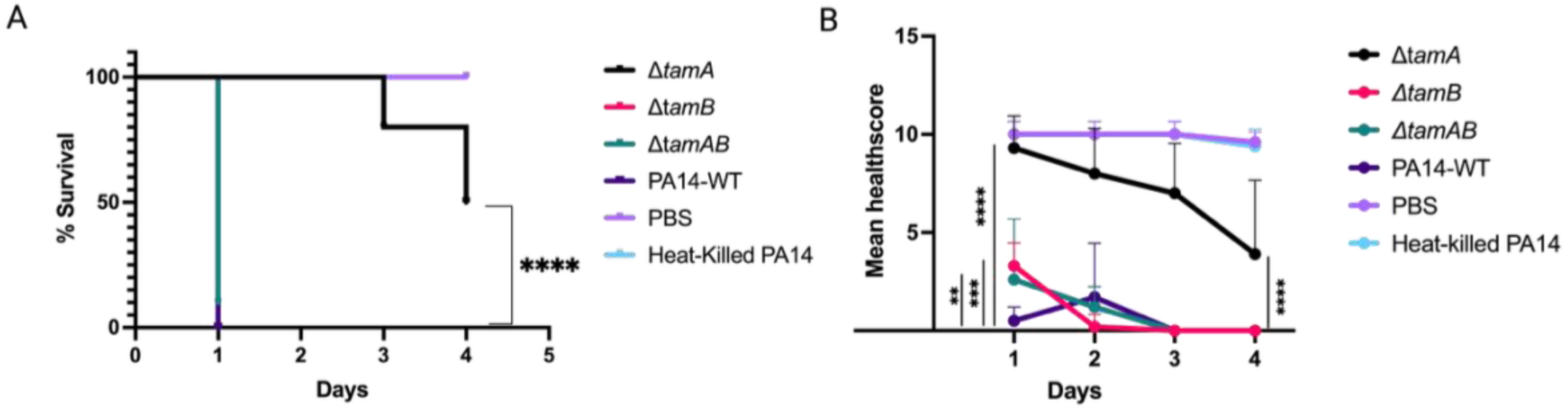
*tamA* knockout is less pathogenic. (A) Survival curve of *Galleria mellonella* post infection. The differences in the survival curves of *ΔtamA* and wildtype are significant using the Mantel-Cox test with ****p <0.0001. (B) Health score of *Galleria mellonella* post infection. The health score of n=10 was calculated daily for four days. Health scores of all TAM mutants and the wildtype for the first day are displayed as follows; Δ*tamA* ****p<0.0001; Δ*tamB* ***p<0.0002; Δ*tamAB* **p< 0.01 shown by the black vertical lines with asterisks). Health score of Δ*tamA-*infected larvae on day 4 shown by black vertical line with asterisks ****p<0.0001. Statistical analysis obtained by Two-way ANOVA with Dunnett’s post-test. 10 CFU of mutants or WT were injected and PBS and heat-killed PA14 used as controls. Larvae were incubated at 37 °C for four days, recording their survival and health score daily in accordance with the *Galleria* scoring system of Serano et al, 2023.

## DISCUSSION

The TAM consist of an integral membrane protein TamA connected by its three POTRA domains to an AsmA-like protein TamB, which spans the periplasm to the inner membrane. The TAM has been demonstrated to play important roles in the assembly of several β-barrel proteins into the OM. (Selkrig et al., 2012; Shen et al., 2014; Stubenrauch et al., 2022; Bamert et al., 2017; Stubenrauch et al., 2016; Josts et al., 2017).

Studies establishing involvement of the TAM in virulence of the opportunistic pathogen *P. aeruginosa* are lacking and in fact only one study has focused on *P. aeruginosa* as a model organism to investigate the involvement of the TAM in phospholipid transport (Sposato et al., 2024). Given that the outer membrane of *P. aeruginosa* contributes greatly to its virulence and overall success as a pathogen (Ozer et al., 2021; Sabnis et al., 2021), compounded with the increasingly needed alternative drugs for treatment of *P. aeruginosa* infections due to antimicrobial resistance, we decided to investigate the TAM as a potential novel drug target.

We assessed the growth potential of *tam* mutants and WT and did not find significant differences between their growth profiles in LB or minimal media, except for Δ*tamA* which had a slightly faster growth rate. Bacteria use a significant amount of their energy to synthesize flagella and drive motility (Berry & Armitage, 1999). Since there is reduced motility exhibited in Δ*tamA* strains, it is possible that energy that would have been used for motility is being directed towards faster growth.

We then decided to investigate the relative fitness of the knockout mutants in a competitive environment by co-culturing each mutant with the WT. We found the mutants were outcompeted by the WT. The inability of *tam* mutants to compete for nutrients equally in a co-culture growth environment with the wildtype could be due to the low level of porins and TonB-dependent receptors present both in Δ*tamA* and Δ*tamB* OM proteome (Table 1). Porins and TonB-dependent receptors facilitate the uptake of nutrients; their reduction in the mutants would result in inefficient competition for nutrients (Chevalier et al., 2017). In a monoculture, however, the reduced transport capacity of mutants does not affect growth because of the lack of competition: the mutants are able to acquire sufficient nutrients for normal growth when the WT is not present, which can absorb limiting nutrients more efficiently.

The OM proteomes of the Δ*tamA* and Δ*tamB* are highly different from the WT. By contrast, the Δ*tamAB* mutant presented with a similar profile to WT where differences observed were not significant. (Figure 2 A, B, C; Supplementary Figure 3). TAM function seems to be limited when there is an absence of either of its subunits. It seems the TAM complex only functions optimally if the two components are present in stoichiometric amounts. This possibly explains why the complementation is often only partial: there is an excess of one component, and consequently the TAM is not functioning optimally. By contrast, deleting the whole *tam* operon had much less of an effect, suggesting that complete absence of the TAM can be compensated for by other cellular machineries, such as the BAM or other AsmA proteins.

As there was mostly a reduction of structural components of flagella (Supplementary Table 4) in the OM proteome, we found that only the *tamA* mutant impacted on swimming motility. Unlike our results, motility was significantly impaired in both *ΔtamA* and *ΔtamB* mutants of *Edwardsiella tarda* (Li et al., 2020). The reduction of flagellar components would have been an explanation for attenuated biofilm formation ability among *tam* mutants; however, the loss of flagellar proteins was not reflected among all the mutants. SPIA and KEGG analysis of the transcriptomic data showed that biofilm formation pathways and quorum sensing pathways were inhibited in all mutants elucidating the reduction in biofilm-forming abilities of the mutants.

With respect to the sulfur relay system (Ikeuchi et al., 2006) detected by SPIA (supplementary Table 5), TusA of the system has been implicated in biofilm formation in *P. aeruginosa* (Filiatrault et al., 2013).When TusBCDE is inhibited, TusE cannot stimulate sulfur transfer from TusA to TusD, essentially disrupting the sulfur relay system.

Channel proteins of the TolC family form the outer membrane component of tripartite efflux pumps. Structural analyses demonstrate that the efflux protein TolC in Gram-negative bacteria functions as a channel for antibiotic removal, influencing bacterial susceptibility and virulence. (Kantarcioglu et al., 2024). Deletion of *tolC* in *E. coli* resulted in increased susceptibility to macrolides, tetracycline and quinolones (Hou et al., 2014). TolC is downregulated 6-fold in both single mutants suggesting a potentially compromised TolC-mediated export of toxic compounds. We would therefore predict that these mutants will have reduced resistance to harmful compounds due to the impaired ability to expel toxic substances like antibiotics and detergents. Some significant differences were observed for detergents and chelators, suggesting that the OM integrity and the ability of the mutants to remove toxic compounds is lowered (Figure 3C).

However, the antibiotics tested against the mutants either showed a reduced effect or no difference compared with the WT (Figure 3 A; supplementary Figure 3). This is in contrast to the increased OM permeability demonstrated by the mutants in the NPN assay, suggesting that the TAM does play an important role in maintaining the barrier of the outer membrane as its removal increases membrane permeability just as previously demonstrated in Δ*tamB* (Ruiz et al., 2021).

Consequently, if drugs are formulated to target the TAM, it should be noted not to use these drugs in combination with the antibiotics that illicit reduced susceptibilities in the TAM knockouts. Other drug/antibiotic combinations may need to be tested.

There is an increase in the production of BamB and BamC potentially due to these lipoproteins assuming a greater role in OMP translocation and insertion in the absence of *tamA*. Deletion of *bamB* has been shown to not only effect changes to the OM permeability (Rollauer et al., 2015), but also to reduce pathogenicity and sensitize bacteria to antibiotics (Fardini et al., 2007).

Similarly, lack of BamC affects OM permeability and sensitizes bacteria to environmental stresses (Onufryk et al., 2005). With the increase of these proteins in the absence of *tamA*, it could be inferred that maintenance of the OM barrier is shifted towards BAM components. We also noticed an effect in LptD expression in the single knockouts which might mean there is an effect on lipopolysaccharide, but this requires further investigation.

To ascertain what these phenotypes might mean in vivo, and to answer our question regarding the TAM’s suitability as a drug target, we infected *Galleria mellonella* larvae and found out the Δ*tamA* mutant is significantly less pathogenic as it had the best health outcome post infection as well as reduced mortality compared to the wild type (Figure 5). This agrees with the findings of Jung et al., (2021) who showed that Δ*tamA* mutants of *Klebsiella pneumoniae* were compromised in their ability to colonize the mouse intestine and easily removed by the host immune system. Similarly, (Selkrig et al,2012) showed that Δ*tamA* or Δ*tamB* mutants of *Citrobacter rodentium* were defective in the colonization of mice. In the same study, it was shown that *Salmonella enterica* SL1344 *tam* mutants became sensitive to human serum.

Although larvae infected with Δ*tamB* died within 24 hours, they showed fewer signs of melanization thus had a better health score, which suggest some level of attenuation of virulence. However, larvae infected with the *tamA* knockout displayed increased survival. This informed our preference of *tamA* as a candidate for drug development.

Altogether, our results point to the importance of the TAM in OMP biogenesis in *P. aeruginosa*. In addition, the negative impact on motility in the absence of the *tamA* gene is due to a reduction in flagellar assembly. Also, TAM mutants have a reduced biofilm forming ability and perform poorly in a competition environment. Further investigation is required to understand why, despite there being an increase in OM in permeability in the TAM mutants, their susceptibility to antibiotics is unchanged or lower. These findings together lead us to propose the Δ*tamA* is a potential drug target against *P. aeruginosa*.

The OM of Gram-negative bacteria poses a major hurdle against the discovery of new antibiotics aimed at treating infections caused by multidrug-resistant bacteria. OM-associated proteins are therefore attractive novel targets for drugs. With advancement with the novel drugs targeting the BamA, for example darobactin (Kaur et al., 2021) and monoclonal antibodies (Storek et al., 2018), similar approaches could be used towards novel drugs against TamA.

## ETHICAL APPROVAL

There is no ethical approval required for the use of *G. mellonella* in the U.K.

## AUTHOR CONTRIBUTIONS

Conceptualization, J.C.L., methodology, J.C.L., D.B., L.H.,; validation, J.CL., and R.D.,; formal analysis, R.D., L.H. and D.B.; investigation, R.D., ; resources, J.C.L .,; data curation, R.D.; writing—original draft preparation R.D,; writing—review and editing, J.C.L., R.D., L.H., DB.; visualization, R.D. and L.H; project administration, J.C.L; funding acquisition, J.C.L., and R.D.

## ACKNOWLEDGEMENTS

We thank Sandra Schwarz (University of Tuebingen, Germany) for gifting us the vector pEXG2 and the strain *P. aeruginosa* PA14.

pSRKGm plasmid was gifted by Clay Fuquah, Indiana University Bloomington, USA

pCMT-flp was a gift from Mark Liles (Addgene plasmid # 67274; http://n2t.net/addgene:67274; RRID: Addgene_67274) and well appreciated.

pFRT-Tet129 was a gift from Doug Mortlock (Addgene plasmid #33359; http://n2t.net/addgene:33359; RRID: Addgene_33359).

pCMT-flp was a gift from Mark Liles (Addgene plasmid # 67274; http://n2t.net/addgene:67274; RRID: Addgene_67274).

pFlp-Gm was a gift from Philip Poole (Addgene plasmid # 222335 ; http://n2t.net/addgene:222335 ; RRID:Addgene_222335).

