## Supplementary material for "The Omp85 family protein, TamA, exhibits characteristics of a suitable drug target against *Pseudomonas aeruginosa*"

### SUPPLEMENTARY MATERIALS

#### Supplementary Table 1: List of primers used in this study

| Primer | Sequence | Comments |
|---|---|---|
| TamAUP_FR | CTCGAGCCCGGGGATCCTAAATGACCATGCAACCCTTCGT | For amplifying fragment upstream of *tamA* |
| TamAUP_Rev | GCCGAATTCCAGCACACTCATTCAGGTCGCCGCCGG | For amplifying fragment upstream of *tamA* |
| TamADS_FR | AGCCGAATTCTGCAGATTGAGTGAAGGCGCTGAAGATC | For amplifying fragment downstream of *tamA* |
| TamADS_Rev | CTGCAGGTCGACTCTAGAGGAATAGCGCCGGCTGG | For amplifying fragment downstream of *tamA* |
| TamAcomp_FR | CACAGGAAACAGCATATGATCACCAGTTGCCGATGCTC | For amplifying tamA locus for insertion into pSRK-Gm |
| TamAcomp_Rev | TTCTAGAGCGGCCGCCAAGTTGCGTACCGTCCTGCG | For amplifying tamA locus for insertion into pSRK-Gm |
| TamBUP_FR | CTCGAGCCCGGGGATCCAGGCCGACGGCGCGCG | For amplifying fragment upstream of *tamB* |
| TamBUP_Rev | GCCGAATTCCAGCACACTCACTCACAGTTCTGGCCCC | For amplifying fragment upstream of *tamB* |
| TamBDS_FR | AAGCCGAATTCTGCAGATTGACGGATGCGCTGATACGGG | For amplifying fragment downstream of *tamB* |
| TamBDS_Rev | CTTCTGCAGGTCGACTCTAGAGGCGATCCGCCCGGAGT | For amplifying fragment downstream of *tamB* |
| pEXG2_FR | TCTAGAGTCGACCTGCAGAAG | For amplifying pEXG2 backbone |
| pEXG2_Rev | GGATCCCCGGGCTCGAG | For amplifying pEXG2 backbone |
| FRT-TET_FR | TGTGCTGGAATTCGGCTTGAAGTTC | For amplifying FRT-TET from pFRT-Tet 129 |
| FRT-TET_Rev | ATCTGCAGAATTCGGCTTGAAGTT | For amplifying FRT-TET from pFRT-Tet 129 |
| pSRK_FR | TGGCGGCCGCTCTAGAAC | For amplifying pSRK-Gm back bone |
| pSRK_Rev | ATGCTGTTTCCTGTGTGAAATT | For amplifying pSRK-Gm back bone |
| pCMTf | AAC CAG GCG TTT AAG GGC ACC AA | For inserting the mSF-ts1 ori into pCMT |
| pCMTr | GCT ACG CCT GAA TAA GTG ATA ATA AGC | For inserting the mSF-ts1 ori into pCMT |
| mSF-pCMTf | CAC TTA TTC AGG CGT AGC CTG CAG AAA GGC AGG CCG GG | for amplifying the mSF-ts1 promoter for insertion into pCMT |
| mSF-pCMTr | G CCC TTA AAC GCC TGG TT CCT CTC AGG CGC CGC TG | |
| pCMT-GmCassF | ACGCGATGGATATGTTCT CAG AGC GCT TTT GAA GCT GAT | for amplifying the Gm cassette for insertion into pCMT |
| pCMT-GmCassR | CAGCCCCATACGATATAA GAT CCC CTG ATT CCC TTT GTC | |
| pCMTf3 | TTATATCGTATGGGGCTGACTTC | for amplifying pCMT to switch the antibiotic cassettes |
| pCMTr3 | AGA ACA TAT CCA TCG CGT CCG | |
| Gm-araBADf | GAA TCA GGG GAT CTT A GAG CTG CAT GTG TCA GAG G | for amplifying the $P_{BAD}$-*araC* insert |
| CMT-araBADr | GAT GGT GGT GAT GAT GCA T GGT TAA TTC CTC CTG TTA GTA | |
| GmF | TAAGATCCCCTGATTCCCTTTG | for amplifying the vector to insert $P_{BAD}$-*araC* |
| pCMTr | ATG CAT CAT CAC CAC CAT CAC | |

Supplementary table 2. List of all differentially expressed proteins (unfiltered)in *ΔtamA* proteome which are significant

| Uniprot Id | Protein Description | LogFC ΔtamA vs PA14 |
|---|---|---|
| A0A077JXD3_PSEAI | Fucose-binding lectin | -7.865929338 |
| A0A0H2ZJM2_PSEAB | alcohol dehydrogenase | -6.907702836 |
| A0A0H2ZH89_PSEAB | Uncharacterized protein | -6.711833929 |
| PHZF_PSEAE | Trans-2,3-dihydro-3-hydroxyanthranilate isomerase | -6.69507018 |
| ACNA_PSEAE | Aconitate hydratase A | -6.661768621 |
| A0A0H2ZE94_PSEAB | Alkyl hydroperoxide reductase | -6.657786159 |
| PNP_PSEAB | Polyribonucleotide nucleotidyltransferase | -6.413638591 |
| CATA_PSEAE | Catalase A | -6.410061983 |
| A0A7M2ZZQ8_PSEAI | Channel protein TolC | -6.287048852 |
| A0A0H2Z9D0_PSEAB | oxoglutarate dehydrogenase (succinyl-transferring) | -6.268308894 |
| A0A0H2ZG18_PSEAB | Putative nonribosomal peptide synthetase | -6.250094362 |
| A0A509JM65_PSEAI | NAD-glutamate dehydrogenase | -6.198699009 |
| ODO2_PSEAE | Dihydrolipoyllysine-residue succinyltransferase component of 2-oxoglutarate dehydrogenase complex | -6.117983608 |
| ILVC_PSEAB | Ketol-acid reductoisomerase (NADP(+)) | -6.079632011 |
| A0A077SYW5_PSEAI | Cyanide-insensitive cytochrome bd quinol oxidase subunit I | -5.996137321 |
| SYV_PSEAE | Valine--tRNA ligase | -5.834216731 |
| PURA_PSEAB | Adenylosuccinate synthetase | -5.826351974 |
| A0A0H2Z6K8_PSEAB | 2,3-dehydroadipyl-CoA hydratase | -5.606538874 |
| RISB_PSEAB | 6,7-dimethyl-8-ribityllumazine synthase | -5.583834168 |
| RL10_PSEAB | Large ribosomal subunit protein uL10 | -5.53846464 |
| A0A0H2ZFE1_PSEAB | Universal stress protein | -5.505684751 |
| ALGR_PSEAE | Positive alginate biosynthesis regulatory protein | -5.495513944 |
| A0A0H2Z834_PSEAB | Flagellar hook-associated protein 2 | -5.475610636 |
| ARCA_PSEAE | Arginine deiminase | -5.458409552 |
| RL31_PSEAB | Large ribosomal subunit protein bL31 | -5.422771312 |
| A0A0H2ZDT9_PSEAB | Chemotaxis protein CheV | -5.362046911 |
| A0A0H2ZDU5_PSEAB | Adenylate cyclase | -5.337051737 |
| EFP_PSEAB | Elongation factor P | -5.332813598 |
| RL16_PSEAB | Large ribosomal subunit protein uL16 | -5.273408281 |
| SYP_PSEAB | Proline--tRNA ligase | -5.25084362 |
| A0A0H2ZCR1_PSEAB | R body protein RebB-like protein | -5.23645196 |
| A0A072ZNQ8_PSEAI | Glycine zipper 2TM domain-containing protein | -5.179865506 |
| RL32_PSEAB | Large ribosomal subunit protein bL32 | -5.148210906 |
|  | Porin | -5.122999336 |
| A0A072ZBY7_PSEAI | TolC family protein | -5.117572791 |
| MQO1_PSEAE | Probable malate:quinone oxidoreductase 1 | -5.112669247 |
| SUCD_PSEAE | Succinate--CoA ligase [ADP-forming] subunit alpha | -5.080746053 |
| EFTS_PSEAB | Elongation factor Ts | -4.978803416 |
| CH60_PSEAB | Chaperonin GroEL | -4.965385073 |
| A0A0H2ZGG4_PSEAB | Putative lipoprotein | -4.953177921 |
| A0A071KW46_PSEAI | LemA family protein | -4.943843523 |
| A0A0F7QY13_PSEAI | TonB-dependent receptor | -4.86532359 |
| MURA_PSEAB | UDP-N-acetylglucosamine 1-carboxyvinyltransferase | -4.836392269 |
| FTNA_PSEAE | Bacterial ferritin | -4.814124726 |
| A0A0H2ZBY5_PSEAB | DUF883 domain-containing protein | -4.80517573 |
| AROC_PSEAB | Chorismate synthase | -4.755118918 |
| A0A0H2Z9V8_PSEAB | Putative universal stress protein | -4.665669955 |
| RS3_PSEAB | Small ribosomal subunit protein uS3 | -4.659004462 |

| | | |
|---|---|---|
| A0A367MEB1_PSEAI | Ferrichrome-iron receptor | -4.614810069 |
| A0A0H2ZI23_PSEAB | Transcription termination/antitermination protein NusA | -4.602545961 |
| SECY_PSEAE | Protein translocase subunit SecY | -4.577720346 |
| A0A0H2Z8M2_PSEAB | Small-conductance mechanosensitive channel | -4.57341222 |
| RRAAH_PSEAB | Putative 4-hydroxy-4-methyl-2-oxoglutarate aldolase | -4.562684972 |
| VFR_PSEAE | cAMP-activated global transcriptional regulator Vfr | -4.559808048 |
| LAP_PSEAB | Aminopeptidase | -4.547764704 |
| A0A0H2ZC12_PSEAB | Autotransporter domain-containing protein | -4.545814389 |
| PYRG_PSEAB | CTP synthase | -4.492035666 |
| A0A0H2Z999_PSEAB | Putative outer membrane receptor protein | -4.482399522 |
| A0A6A9KBX7_PSEAI | TonB-dependent siderophore receptor | -4.465705573 |
| OTCC_PSEAE | Ornithine carbamoyltransferase, catabolic | -4.45061828 |
| IDH_PSEAB | Isocitrate dehydrogenase [NADP] | -4.443132552 |
| A0A0H2Z9D1_PSEAB | Transcriptional regulator CysB | -4.383737076 |
| RL28_PSEAB | Large ribosomal subunit protein bL28 | -4.35198133 |
| A0A0H2ZDI1_PSEAB | Putative zinc-binding dehydrogenase | -4.350925315 |
| A0A0H2ZK67_PSEAB | ATP-dependent RNA helicase RhlE | -4.225342229 |
| A0A0H2ZLD6_PSEAB | Nitrite reductase | -4.223383052 |
| PHOP_PSEAE | Two-component response regulator PhoP | -4.215820831 |
| A0A0H2ZFL2_PSEAB | Putative outer membrane protein | -4.209489762 |
| A0A0H2ZI19_PSEAB | DUF748 domain-containing protein | -4.204256511 |
| A0A0H2ZKH1_PSEAB | Efflux RND transporter periplasmic adaptor subunit | -4.161469157 |
| FLEQ_PSEAE | Transcriptional regulator FleQ | -4.152447594 |
| A0A0H2Z762_PSEAB | Exonuclease VII large subunit C-terminal domain-containing protein | -4.115454865 |
| NUOH_PSEAB | NADH-quinone oxidoreductase subunit H | -4.105216058 |
| PILY1_PSEAB | Type IV pilus biogenesis factor PilY1 | -4.058229298 |
| SSB_PSEAE | Single-stranded DNA-binding protein | -3.989148826 |
| HEM2_PSEAE | Delta-aminolevulinic acid dehydratase | -3.938687862 |
| RS14_PSEAB | Small ribosomal subunit protein uS14 | -3.853930137 |
| A0A0H2ZKT8_PSEAB | Transketolase | -3.849767603 |
| A0A0H2ZL81_PSEAB | Iron-sulfur cluster carrier protein | -3.717451582 |
| A0A0H2ZKE8_PSEAB | C-type cytochrome | -3.632774412 |
| RL30_PSEAB | Large ribosomal subunit protein uL30 | -3.565629276 |
| A0A0H2ZIJ0_PSEAB | Carbamoyltransferase | -3.421879316 |
| RS20_PSEAB | Small ribosomal subunit protein bS20 | -3.360146968 |
| A0A0H2ZGA2_PSEAB | RND efflux membrane fusion protein | -3.313169301 |
| HCP1_PSEAE | Protein hcp1 | -3.264978678 |
| FTSA_PSEAE | Cell division protein FtsA | -3.215474622 |
| A0A0H2ZHF8_PSEAB | Putative outer membrane protein | -2.439195285 |
| A0A0H2Z8K3_PSEAB | Sn-glycerol-3-phosphate transporter | -1.306013263 |
| A0A6A9JVT0_PSEAI | DUF533 domain-containing protein | 1.705979976 |
| A0A0H2ZDL5_PSEAB | Peptidyl-prolyl cis-trans isomerase | 1.857560154 |
| A0A0H2ZES8_PSEAB | Putative lipoprotein | 1.945347733 |
| A0A0H2ZHP1_PSEAB | Paraquat-inducible protein B-like protein | 2.004496465 |
| ATPA_PSEAB | ATP synthase subunit alpha | 2.187650967 |
| A0A3M5EB07_PSEAI | BON domain-containing protein | 2.308352261 |
| A0A0H2ZJN1_PSEAB | Outer membrane assembly lipoprotein YfiO | 2.421652005 |
| A0A0H2Z890_PSEAB | Putative lipoprotein | 2.451576231 |
| A0A0H2ZDH6_PSEAB | General secretion pathway protein D | 2.669518389 |
| YFIB_PSEAE | Outer-membrane lipoprotein YfiB | 3.03583481 |
| A0A0H2ZIC6_PSEAB | OsmE family transcriptional regulator | 3.138522732 |
| A0A069QEN7_PSEAI | Chromosome partitioning protein ParA | 3.700437505 |
| BAMB_PSEAE | Outer membrane protein assembly factor BamB | 4.282713113 |

Supplementary table 3. List of all differentially expressed proteins (unfiltered)in *ΔtamB* proteome which are significant

| UniProt Id | Protein Description | LogFC *ΔtamB* vs PA14 |
|---|---|---|
| A0A0H2ZG05_PSEAB | Putative porin | -7.824455707 |
| ALGP_PSEAE | Transcriptional regulatory protein AlgP | -6.889544021 |
| ACNA_PSEAE | Aconitate hydratase A | -6.661768621 |
| A0A643IRS3_PSEAI (Porin) | Porin | -6.642985962 |
| A0A0H2ZA96_PSEAB | Transporter | -6.511634235 |
| PNP_PSEAB | Polyribonucleotide nucleotidyltransferase | -6.413638591 |
| CATA_PSEAE | Catalase A | -6.410061983 |
| A0A0H2ZE85_PSEAB | Fucose-binding lectin | -6.33127282 |
| A0A0H2Z917_PSEAB 9 | Putative purine-binding chemotaxis protein CheW | -6.326043707 |
| A0A7M2ZZQ8_PSEAI | Channel protein TolC | -6.287048852 |
| A0A0H2Z9D0_PSEAB | oxoglutarate dehydrogenase (succinyl-transferring) | -6.268308894 |
| A0A0H2ZC71_PSEAB | Putative secretion system protein | -6.123551896 |
| ODO2_PSEAE | Dihydrolipoyllysine-residue succinyltransferase component of 2-oxoglutarate dehydrogenase complex | -6.117983608 |
| ILVC_PSEAB | Ketol-acid reductoisomerase (NADP(+)) | -6.079632011 |
| NUOCD_PSEAB | NADH-quinone oxidoreductase subunit C/D | -5.786004306 |
| DNAK_PSEAB | Chaperone protein DnaK | -5.667103455 |
| A0A0H2ZEU8_PSEAB | Alanine/glycine:cation symporter family protein | -5.585263042 |
| A0A0H2ZFH3_PSEAB | serine-type D-Ala-D-Ala carboxypeptidase | -5.538248955 |
| A0A0H2ZE94_PSEAB | Alkyl hydroperoxide reductase | -5.492724516 |
| RRAAH_PSEAB | Putative 4-hydroxy-4-methyl-2-oxoglutarate aldolase | -5.477959355 |
| A0A0H2Z834_PSEAB | Flagellar hook-associated protein 2 | -5.475610636 |
| ARCA_PSEAE | Arginine deiminase | -5.458409552 |
| RL4_PSEAB | Large ribosomal subunit protein uL4 | -5.374887604 |
| A0A0H2ZDT9_PSEAB | Chemotaxis protein CheV | -5.362046911 |
| RL24_PSEAB | Large ribosomal subunit protein uL24 | -5.348195756 |
| A0A509JL21_PSEAI | Adenylate cyclase | -5.337051737 |
| SYP_PSEAB | Proline--tRNA ligase | -5.25084362 |
| A0A0H2ZCR1_PSEAB | R body protein RebB-like protein | -5.23645196 |
| RL32_PSEAB | Large ribosomal subunit protein bL32 | -5.148210906 |
| A6V673_PSEP7 | Porin | -5.122999336 |
| A0A0S2TT89_PSEAI | TolC family protein | -5.117572791 |
| MQO1_PSEAE | Probable malate:quinone oxidoreductase 1 | -5.112669247 |
| SUCD_PSEAE | Succinate--CoA ligase [ADP-forming] subunit alpha | -5.080746053 |
| SYV_PSEAE | Valine--tRNA ligase | -5.027604472 |
| ATPF_PSEAB | ATP synthase subunit b | -4.989355619 |

| | | |
|---|---|---|
| EFTS_PSEAB | Elongation factor Ts | -4.978803416 |
| A0A0H2ZCQ2_PSEAB | Putative TonB-dependent receptor | -4.967871431 |
| A0A0H2ZGG4_PSEAB | Putative exported lipoprotein | -4.953177921 |
| A0A0H2ZKG2_PSEAB | LemA family protein | -4.943843523 |
| A0A0H2ZIC7_PSEAB | Protein HflC | -4.923096366 |
| A0A0H2Z6U8_PSEAB | TonB-dependent receptor | -4.86532359 |
| LLDD_PSEAB | L-lactate dehydrogenase | -4.814039313 |
| A0A0H2ZBY5_PSEAB | DUF883 domain-containing protein | -4.80517573 |
| AROC_PSEAB | Chorismate synthase | -4.755118918 |
| RS3_PSEAB | Small ribosomal subunit protein uS3 | -4.659004462 |
| A0A509JN15_PSEAI | Glucose/quinate/shikimate family membrane-bound PQQ-dependent dehydrogenase | -4.617284117 |
| A0A3M5DNP1_PSEAI; | Ferrichrome-iron receptor | -4.614810069 |
| SECY_PSEAE | Protein translocase subunit SecY | -4.577720346 |
| A0A0H2Z8M2_PSEAB | Small-conductance mechanosensitive channel | -4.57341222 |
| VFR_PSEAE | cAMP-activated global transcriptional regulator Vfr | -4.559808048 |
| LAP_PSEAB | Aminopeptidase | -4.547764704 |
| PYRG_PSEAB | CTP synthase | -4.492035666 |
| A0A0H2Z999_PSEAB | Putative outer membrane receptor protein | -4.482399522 |
| A0A6A9KBX7_PSEAI | TonB-dependent siderophore receptor | -4.465705573 |
| OTCC_PSEAE | Ornithine carbamoyltransferase, catabolic | -4.45061828 |
| A0A0H2ZJ18_PSEAB | Cytochrome c5 | -4.406042472 |
| A0A0H2Z9D1_PSEAB | Transcriptional regulator CysB | -4.383737076 |
| A0A0H2ZDI1_PSEAB | Putative zinc-binding dehydrogenase | -4.350925315 |
| PCTB_PSEAE | Methyl-accepting chemotaxis protein PctB | -4.245178071 |
| FTSZ_PSEAE | Cell division protein FtsZ | -4.220989016 |
| A0A0H2ZFL2_PSEAB | Putative outer membrane protein | -4.209489762 |
| A0A0H2Z762_PSEAB | Exonuclease VII large subunit C-terminal domain-containing protein | -4.115454865 |
| PILY1_PSEAB | Type IV pilus biogenesis factor PilY1 | -4.058229298 |
| A0A072ZIB6_PSEAI | Cytochrome c oxidase accessory protein CcoG | -4.057256562 |
| A0A0H2ZLD6_PSEAB | Nitrite reductase | -4.021409038 |
| SSB_PSEAE | Single-stranded DNA-binding protein | -3.989148826 |
| A0A0H2ZG66_PSEAB | Putative iron-sulfur cluster-binding protein | -3.98879022 |
| MURA_PSEAB | UDP-N-acetylglucosamine 1-carboxyvinyltransferase | -3.973915905 |
| A0A0H2ZKM8_PSEAB | Putative hydroxamate-type ferrisiderophore receptor | -3.968357158 |
| A0A0H2Z825_PSEAB | Flagellar basal-body rod protein FlgG | -3.94962816 |
| HEM2_PSEAE | Delta-aminolevulinic acid dehydratase | -3.938687862 |
| A0A071KU93_PSEAI | C-type cytochrome | -3.632774412 |
| ATPD_PSEAB | ATP synthase subunit delta | -3.593188756 |
| ETFD_PSEAE | Electron transfer flavoprotein-ubiquinone oxidoreductase | -3.540848232 |

| | | |
|---|---|---|
| DNAJ_PSEAB | Chaperone protein DnaJ | -3.481737828 |
| A0A0H2ZIJ0_PSEAB | Carbamoyltransferase | -3.421879316 |
| A0A0H2ZGA2_PSEAB | RND efflux membrane fusion protein | -3.313169301 |
| FTSA_PSEAE | Cell division protein FtsA | -3.215474622 |
| A0A0H2ZHF8_PSEAB | Putative outer membrane protein | -2.439195285 |
| A0A367ME26_PSEAI | DUF4823 domain-containing protein | 1.569201846 |
| A0A069QEN7_PSEAI | Chromosome partitioning protein ParA | 1.933965135 |
| BAMB_PSEAE | Outer membrane protein assembly factor BamB | 1.941419858 |
| A0A0H2ZHP1_PSEAB | Paraquat-inducible protein B-like protein | 2.025882742 |
| A6V2I3_PSEA7 | Lipoprotein, putative | 2.027421614 |
| A0A0H2ZDL5_PSEAB | Peptidyl-prolyl cis-trans isomerase | 2.09093722 |
| A0A0H2ZIY0_PSEAB | Dodecin family protein | 4.191385679 |

Supplementary Table 4. List of all differentially expressed flagellar components in *ΔtamA, ΔtamB* and *ΔtamAB OM* proteome .

| UniProt Id | Protein Description | LogFC *ΔtamA* vs PA14 | LogFC *ΔtamB* vs PA14 | LogFC *ΔtamAB* vs PA14 |
|---|---|---|---|---|
| A0A0H2Z834_PSEAB | Flagellar hook-associated protein 2 | -5.475610636 | -5.475610636 | -1.020982816 |
| A0A0H2Z7W1_PSEAB | Flagellar hook protein FlgE | -3.94424859 | -2.279062192 | -0.383820485 |
| FLIF_PSEAE | Flagellar M-ring protein | -3.592361808 | -3.430941691 | 0.022923384 |
| A0A0H2Z825_PSEAB | Flagellar basal-body rod protein FlgG | -2.794433263 | -3.94962816 | -0.563453483 |
| *P72129* | Flagellin (Fragment) | -2.326370961 | -0.384583319 | -0.593177766 |
| *A0A6M5K8Z5* | B-type flagellin (Fragment) | -1.464217103 | -1.464217103 | 3.058340548 |
| A3RJ52_PSEAI | Flagellin | -1.088882125 | 0.572190949 | -0.243446832 |
| FLGH_PSEAB | Flagellar L-ring protein | -0.834778256 | 0.733876705 | 0.197317268 |
| *FLGI_PSEAB* | Flagellar P-ring protein | -0.049199069 | -1.481075555 | -0.197508165 |
| *A0A6M5KA93* | B-type flagellin (Fragment) | 0.905582021 | 3.190088506 | -7.204838758 |

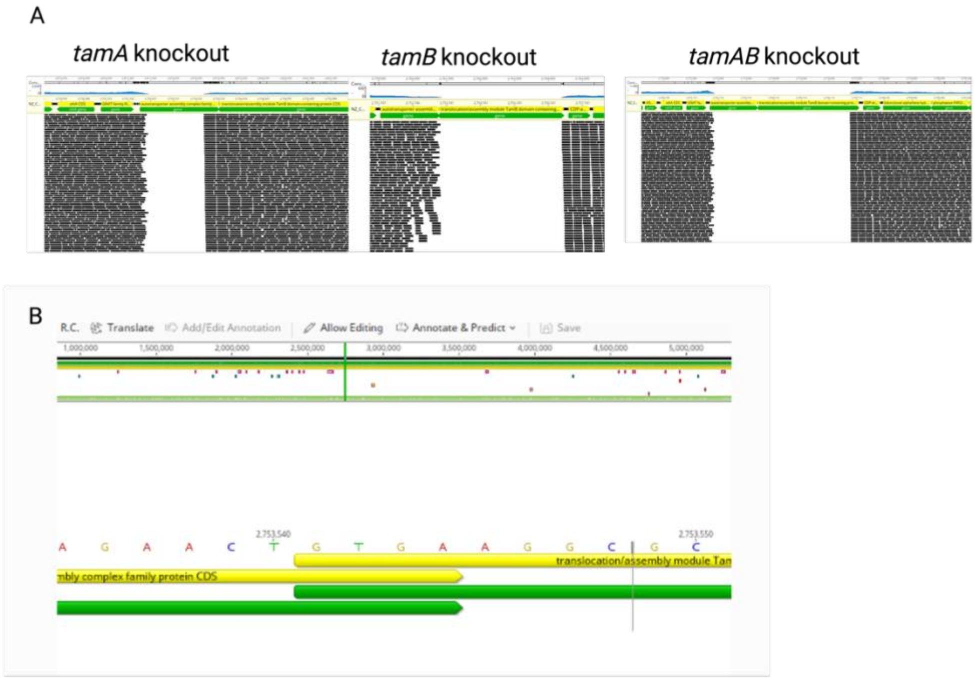


**Supplementary Figure 1.**
**tam mutants and genome**
(A) reads of tam gene knockouts mapped against reference strain *P. aeruginosa PA14*
(B) overlapping stop codon of *tamA* with *tamB*

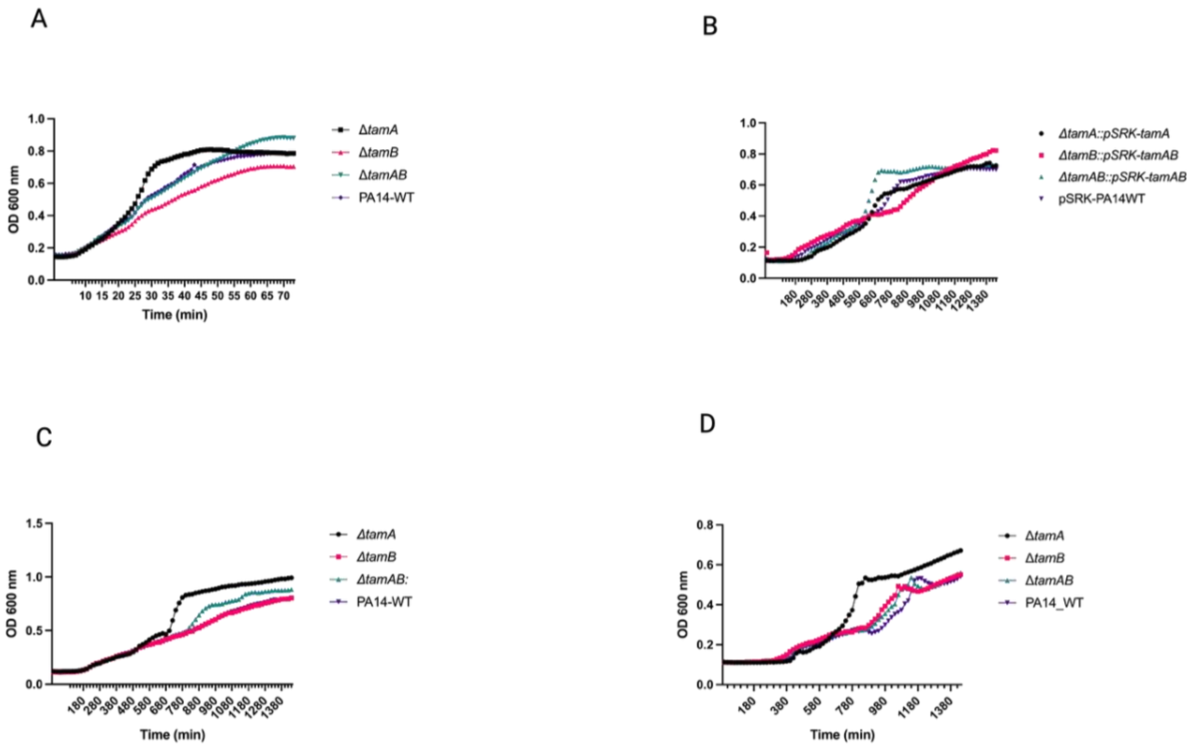


**Supplementary Figure 2.**

**Growth profiles of tam mutants compared to the wildtype**

(A) Growth curve of tetracycline sensitive mutants vs wildtype, (B) Growth curve of complemented TAM mutants vs wildtype containing pSRK background (C), Growth curve of tetracycline resistant TAM mutants compared to the wildtype showing curves represented by mean of four biological replicates. (D). Growth curve of TAM mutants in M9 media

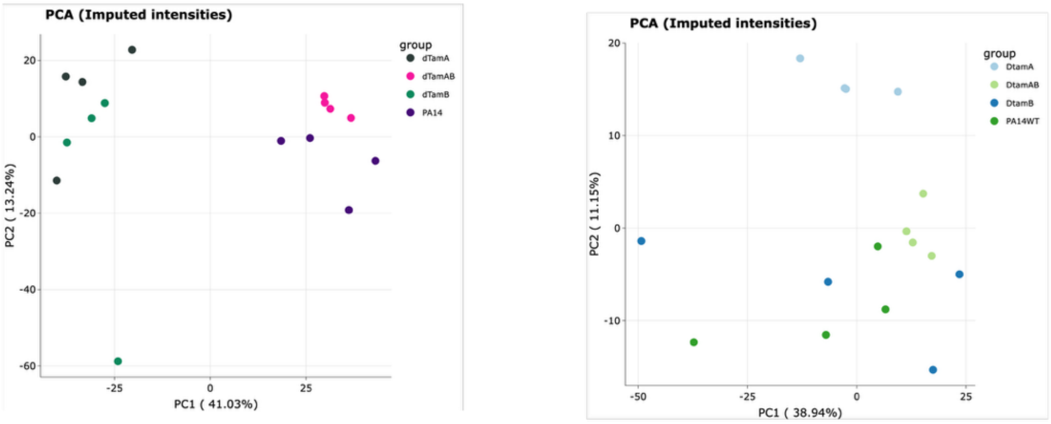


**Supplementary Figure 3**

**Comparative principal component analysis (PCA) of OM and WCP proteome highlights tam mutants vs wildtype separation**

Visualization of the overall variation in protein intensities following imputation using Principal Component Analysis (PCA). (A) PCA of outer membrane proteins (OMPs). The PC1 depicts 41.03% of the variance while PC2 shows 13.24% of the variance in total outlining 54.27% of the total variance for clustering. (B)Whole Cell proteome (WCP) PC1 and PC2 with 38.94% and 11.15% variance respectively. The PCA shows that the *ΔtamA* mutant has a distinct global protein expression pattern, separating strongly from the wildtype and other mutants, especially along PC1. *ΔtamAB* seems to show a somewhat intermediate pattern, while Δ*tamB* alone is more similar to wildtype. This suggests tamA may have a larger impact on the outer membrane proteome than *ΔtamB*.

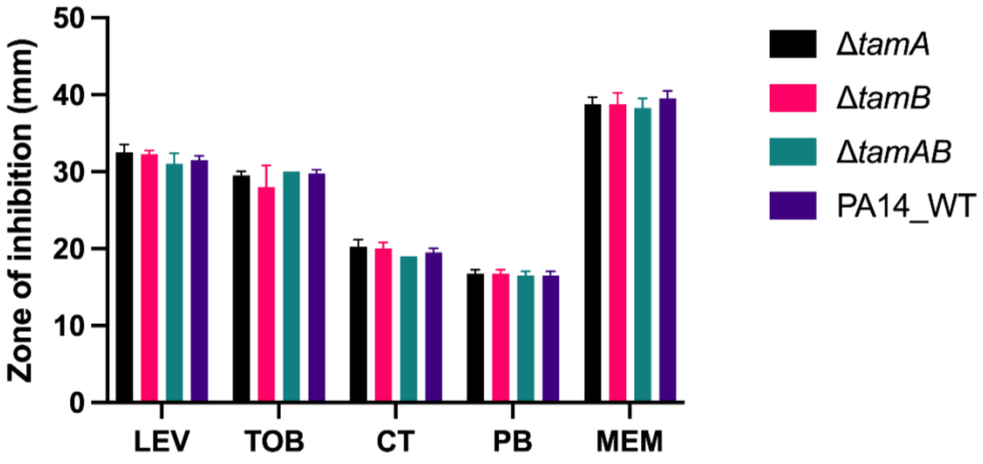


**Supplementary Figure 4**

***tam* mutants and wildtype exhibit comparable susceptibility to Clinically relevant antibiotics**

Antibiotics susceptibility tests for Levofloxacin (LEV), Tobramycin (TOB), Colistin (CT), PolymyxinB (PB) and meropenem (MEM) show no significant differences between mutants and wildtype. 2-way ANOVA was used statistics analysis. Nonsignificant (ns) p values not shown.

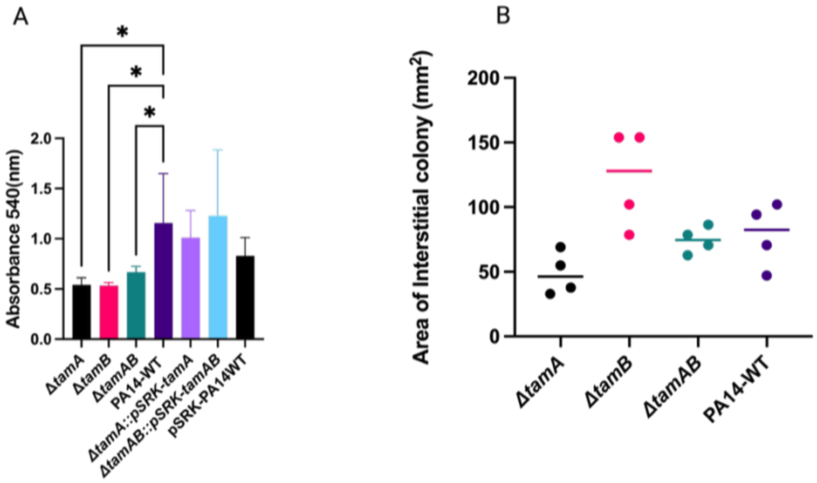


**Supplementary Figure 5.**

**Assessment of Biofilm formation upon complementation and twitching motility of *tam* mutants compared to the WT.**

(A) Complemented strains did not result in total restoration of biofilm-forming ability however there is an increase in biomass compared to the WT.

(B) Twitching of *tam* mutants and wildtype. Twitching motility was performed according to the macroscopic twitching assay of (Turnbull & Whitchurch, 2014).No significant differences observed.

Statical analysis done using One-way Anova. Complemented strains with ns p value not shown.
